# A Comparative Metagenomic Analysis of the Viral Diversity in Faecal Samples from Malnourished and Non-malnourished Children with Gastroenteritis Admitted at Victoria Mxenge Hospital in Durban, KwaZulu-Natal, South Africa

**DOI:** 10.64898/2026.11.24.768824

**Authors:** Marcelle M. Eyong, Milton T. Mogotsi, Hlengiwe Sondlane, Ayodeji E. Ogunbayo, Vivian C. Nwokorogu, Nonkululeko G. Mabaso, Mathew D. Esona, Nathlee Shaik, Kimesh L. Naidoo, Saheed Sabiu, Martin M. Nyaga

## Abstract

Viral gastroenteritis and malnutrition remain major health burdens among children under five in South Africa. Malnourished children are particularly vulnerable to prolonged enteric viral infections, yet their gut virome remains poorly characterised in South Africa. This study compared gut virome diversity among malnourished and non-malnourished children with gastroenteritis, and children without gastroenteritis and malnutrition (healthy controls). This retrospective laboratory-based continuation study analysed 69 faecal samples from age-matched children aged 0– 36 months at Victoria Mxenge Hospital across three cohorts: malnourished children with gastroenteritis, non-malnourished children with gastroenteritis, and healthy controls. Viral metagenomic sequencing was performed using Illumina NextSeq 2000, followed by bioinformatic analyses. Virome profiling identified 39 viral families, 71 genera, and 243 species. Of the total sequenced reads, 60.4% were classified comprising 91.0% mapped to bacteria, 6.4% to viruses, and 2.6% to human host and other microorganisms. Eukaryotic viral reads were highest among non-malnourished children with gastroenteritis (37.2%), followed by healthy controls (32.3%) and malnourished children with gastroenteritis (30.5%). Detected enteric viral families included, *Sedoreoviridae, Caliciviridae*, *Picornaviridae*, *Astroviridae*, and *Adenoviridae,* with *Sedoreoviridae* being predominant. Multiple co-detections of enteric viruses were observed. Significant cohorts differences occurred in viral diversity (*p* = 0.0038) and abundance (*p* = 0.0001), but not in evenness (*p* = 0.146) and richness (*p* = 0.0896). This study revealed differences in faecal virome composition across cohorts and provides baseline genomic data for viral surveillance. The findings highlight the importance of vaccination, nutritional and preventive strategies to curb infant mortality associated with gastroenteritis and malnutrition in South Africa.

## 1.0 Introduction

Gastroenteritis and malnutrition are closely interconnected global public health challenges, particularly among children under five years of age in developing countries (WHO 2024; Hijazi et al., 2022). Diarrhoea, the hallmark clinical manifestation of gastroenteritis, is defined as the passage of three or more loose or watery stools within a 24-hour period (Chukwu et al., 2020). Transmission occurs predominantly through the faecal–oral route via contaminated food and water (Xavier et al., 2020). Globally, gastroenteritis is responsible for approximately 1.7 billion episodes of childhood diarrhoea and an estimated 525,000 deaths annually among children younger than five years (Omatola et al., 2024). It is the third leading cause of death in children 1-59 months of age (WHO 2024). The burden of diarrhoeal disease is greatest in low- and middle-income countries, where malnutrition, inadequate sanitation and hygiene, poor access to healthcare, early weaning, and suboptimal vaccination coverage contribute substantially to morbidity and mortality (Hartman et al., 2023).

Gastroenteritis is caused by a wide range of infectious agents, including viruses, bacteria, protozoa, fungi, and parasites (Xavier et al., 2020). Advances in molecular diagnostic technologies, particularly next-generation sequencing (NGS), have enhanced the detection and characterisation of enteric viruses, which are important aetiological agents of paediatric gastroenteritis. Viruses have been estimated to account for approximately 70% of gastroenteritis cases in children worldwide, rotavirus, norovirus, adenovirus, and astrovirus among the major viral pathogens (Elliott, 2007). The burden of paediatric gastroenteritis is particularly high in sub-Saharan Africa and South-East Asia (Hugho et al., 2023). Nevertheless, approximately 40% of diarrhoeal cases remain of unknown aetiology, suggesting that additional viral and other infectious agents remain to be identified (Xavier et al., 2020).

Before the introduction of the Rotarix vaccine into South Africa’s national childhood immunisation programme in August 2009, rotavirus was responsible for approximately one in four diarrhoea-related hospitalisations among children (Chukwu et al., 2020; Steele et al., 2003). Although the implementation of rotavirus vaccination has substantially reduced the incidence of rotavirus-associated diarrhoea in South Africa and other countries, viral gastroenteritis remains a major cause of childhood illness, highlighting the continued contribution of both established and emerging enteric viruses to paediatric disease (Chukwu et al., 2020; Groome et al., 2014).

Major eukaryotic viruses associated with infant gastroenteritis worldwide include *Rotavirus, Norovirus, Sapovirus*, *Adenovirus,* and *Astrovirus* (Rossouw et al., 2021). Contributing viruses include *Enterovirus, Parechovirus, Kobuvirus, Cosavirus,* and *Cardiovirus* (Xavier et al., 2020). Many of these viruses have also been detected in faecal samples from clinically asymptomatic infants, indicating that viral carriage during early childhood is common (Mogotsi et al., 2020; Yinda et al., 2019). In addition, several plant virus families, including *Alphaflexiviridae, Tombusviridae, Virgaviridae, Solemoviridae, and Geminiviridae* have been identified in the human gut, with their presence thought to reflect dietary and environmental exposure rather than active infection (Blanca et al., 2021). *Siphoviridae, Inoviridae, Myoviridae,* and *Podoviridae,* belonging to the order *Caudovirale*s, together with crAss-like bacteriophages, are among the most abundant bacteriophages inhabiting the human gastrointestinal tract (GIT) (Shkoporov et al., 2019). These bacteriophages constitute a major component of the healthy gut virome and play important roles in shaping bacterial community structure, maintaining microbial homeostasis, and influencing host health.

Malnutrition contributes to approximately 45% of deaths among children under five years globally and remains a major public health concern, particularly in sub-Saharan Africa and South-East Asia, despite declining prevalence worldwide (WHO, 2024; WHO, 2020). It is primarily associated with food insecurity and inadequate nutrient intake, resulting in underweight, stunting, wasting, impaired growth, increased susceptibility to infections, and in severe cases, death (Indiran et al., 2021; Davis et al., 2020). Nutritional status is assessed using anthropometric indices, including height-for-age, weight-for-height, and weight-for-age (WHO, 2011). Globally, an estimated 144 million children were stunted, 47 million wasted, and 38.3 million overweight or obese in 2020 (Clark et al., 2020). In South Africa, the prevalence of stunting, underweight, and wasting among children under five years was estimated at 27.0%, 6.0%, and 2.5%, respectively (May et al., 2021). Childhood malnutrition is influenced by socioeconomic and environmental factors, such as poverty, inadequate feeding practices, poor sanitation, limited access to healthcare, and recurrent infections (Makanjana and Naicker, 2021; Drammeh et al., 2019). Acute malnutrition is associated with increased morbidity, mortality, impaired physical and cognitive development, and greater susceptibility to co-infections (Wali et al., 2019). In children with viral gastroenteritis malnutrition may increase susceptibility to prolonged and recurrent diarrhoeal illness. Rotavirus vaccines may also be less effective in malnourished children than in well-nourished children (Burnett et al., 2021).

The advent of viral metagenomics has greatly advanced the characterization of the human gut virome, enabling the detection of both known and novel viral pathogens (Yang et al., 2022). Recent studies employing metagenomic next-generation sequencing (mNGS) have revealed that the gut, particularly in children under five years of age, harbours a highly diverse and complex viral community (Yang et al., 2022; Santiago-Rodriguez and Hollister, 2020). These investigations have also demonstrated important associations between the gut virome and a range of human diseases (Bai et al., 2022; Wang et al., 2022; Fulci et al., 2021). Despite these advances, the human gut virome remains far less understood than the bacterial microbiome (Blanca et al., 2021). Moreover, the faecal virome of malnourished children under five years of age with gastroenteritis remains poorly characterized in South Africa, despite the high burden of diarrhoeal disease in this vulnerable population (Mafokwane et al., 2023) and the increasing application of mNGS technologies.

Considering the foregoing, this study investigated and compared the faecal virome composition and diversity in malnourished children with gastroenteritis, and non-malnourished children with gastroenteritis, admitted at Victoria Mxenge Hospital, Durban, South Africa, using viral metagenomic next-generation sequencing (mNGS) and bioinformatic analyses. This study provides the first comprehensive baseline data on the gut virome of malnourished children with gastroenteritis in South Africa. The findings may enhance understanding of the viral aetiology of paediatric gastroenteritis and provide evidence to inform future viral surveillance and the development of strategies for preventing and managing viral gastroenteritis and malnutrition.

## 2.0 Materials and Methods

### 2.1. Ethics approvals

This study was reviewed and approved by the Biomedical Research Ethics Committee (BREC) of the University of KwaZulu-Natal under ethics number BREC/000040001/2022 and renewed on 20/03/2025. Approvals were also obtained from the KwaZulu-Natal Department of Health (KZ_202112_002) and the management of Victoria Mxenge Hospital (KE 2/7/16 (11/2022). The study was further reviewed and approved by the Health Sciences Research Ethics Committee (HSREC) under ethics number UFS-HSD2024/2670 and by the Environment and Biosafety Research Ethics Committee (EBREC) under ethics number UFS-ESD2024/0440 of the University of the Free State, Bloemfontein, South Africa.

### 2.2. Study Design and Participants

This was a retrospective, laboratory-based continuation study using archived clinical data and faecal samples from a previous cross-sectional study conducted between January 2023 and October 2024. The study population included malnourished and non-malnourished inpatient or outpatient children with gastroenteritis between 0 to 36 months old at Victoria Mxenge Hospital in Durban, KwaZulu-Natal. The malnourished children suffered from severe acute malnutrition (SAM). According to the World Health Organization (WHO) Anthro software/Anthro Survey Analyser, during sampling, a child was considered to suffer from SAM based on the following criteria: wasting defined as a weight-for-height/length (WHZ/WLZ) Z-score ˂ -3 SD; underweight, defined as a weight-for-age (WAZ) Z-score ˂ -2 SD; stunting, defined as a height-for-age (HAZ) Z-score ˂ -2 SD, mid upper arm circumference (MUAC) was ˂115mm, and the presence of bilateral pitting oedema of both feet. For NAM, WHZ ≥ -2 SD, with MUAC ≥125mm, and no bilateral pitting oedema.

### 2.3. Consent and sample collection

Written consent was obtained from the parents or guardians of the children prior to sample collection after they were given comprehensive information about the research study. Fresh faecal samples were collected per child at a single point in time following defecation in a napkin with the aid of a sterile swab and placed in a 15 mL sterile tube containing 2.5 mL of viral transport medium and properly labelled (date of sample collection and unique sample identification). A total of 71 faecal samples (n= 71) were collected from three age-matched cohorts of inpatient and outpatient children. The first cohort consisted of children with gastroenteritis and malnutrition; a total of 24 samples were collected (n= 24). The second cohort: non-malnourished children with gastroenteritis (n=24), and, lastly, those without gastroenteritis and malnutrition (n=23). One sample was excluded from the cohort consisting of malnourished children with gastroenteritis because of a duplicate sample ID. The second was excluded from the non-malnourished children with gastroenteritis group because the child was 72 months old and therefore did not meet the study’s age eligibility criteria. Consequently, a total of 69 samples were retained for further laboratory analysis, comprising 23 samples from the cohort of malnourished inpatients children with gastroenteritis, 23 samples from the cohort of non-malnourished children with gastroenteritis (22 inpatients and one outpatient), and 23 samples from the cohort of children without gastroenteritis and malnutrition (18 inpatients and five outpatients) (Supplementary Table S1). The samples were stored at -80°C pending laboratory analysis.

### 2.4. Demographic and clinical data

Demographic and clinical metadata, not limited to age, height, weight, gender, breastfeeding or formula, HIV status, and nutritional status of each child, were collected using a well-structured questionnaire. The nutritional status of each child was determined using WHO definitions and anthropometric measurements. The nutritional status was calculated as follows: the weight-for-age Z-score (WAZ ˂ -2 SD) for underweight, weight-for-height Z-score (WHZ ˂ -3 SD) for wasting and height-for-age Z-score (HAZ ˂ -2 SD) for stunting (WHO Anthro for Personal Computers, 2011).

### 2.5. Faecal virome enrichment and nucleic acid extraction

Faecal virome enrichment was performed using a modified version of the Novel Enrichment Technique of Virome (NetoVIR) protocol (Conceição-Neto et al., 2015). Briefly, approximately 100 mg of faecal material was resuspended in 1 mL of sterile phosphate-buffered saline (PBS) (Sigma-Aldrich, St Louis, MO, USA) and homogenised using a Beadbug microtube homogeniser (Benchmark Scientific, Sayreville, NJ, USA) at 3 000 rpm for 1 minute to obtain a 10% (m/v) faecal suspension. This was followed by centrifugation at 13 500 rpm for 3 minutes using Prism microcentrifuge (Labnet, Edison, NJ, USA). A 150 µL volume of the supernatant was retrieved and filtered using a 0.45 µm sterile syringe filter (JLW Supplies, Cape Town, South Africa), and a final volume of 130 µL of the filtrate was collected, to which was added 7 µL of 20x in-house buffer consisting of 1M Tris, 100 mM CaCl_2_ and 30 mM MgCl_2_, pH 8. This was followed by the addition of 2 µL of benzonase (Merck Millipore, Burlington, MA, USA) and 1 µL of micrococcal nuclease (Thermo Fisher Scientific, Meridian Road, Rockford, IL, USA). The tubes were incubated for 2 hours at 37°C in a heating block to digest any extracellular free-floating nucleic acids, and the reaction was stopped by adding 7 µL of 0.5 M ethylenediaminetetraacetic acid (EDTA). Total nucleic materials (DNA and RNA) were extracted using QIAamp Viral RNA Mini Kit 250 (Qiagen, Hilden, Germany), without carrier RNA, following the manufacturer’s instructions.

### 2.6. Whole Transcriptome Amplification (cDNA Synthesis)

Whole transcriptome amplification (cDNA synthesis) was performed using QIAseq Single Cell RNA Library Kit (Qiagen, 180312, Hilden, Germany). The cDNA concentrations were measured on a Qubit 3.0 Fluorometer (Thermo Fisher Scientific) using Invitrogen Qubit ^TM^ 1X dsDNA High Sensitivity Assay Kit (Thermo Fisher Scientific, Q33231, Waltham, MA, United States).

### 2.7. DNA library preparation and next generation sequencing

DNA libraries were prepared on the Hamilton Microlab STAR liquid-handling platform (Hamilton Company, USA) using the Illumina DNA Library Preparation Kit (Illumina, San Diego, California, USA), uniquely indexed with the Illumina DNA/RNA Unique Dual Indexes Set A (Illumina 20091646). The fragment sizes of the DNA libraries were determined on Agilent 2100 Bioanalyzer (Agilent Technologies, Waldbronn, Germany), using Agilent High Sensitivity DNA Kit (5067-4626). The indexed libraries were fluorometrically quantified in ng/µL on a Qubit 3.0 Fluorometer (Thermo Fisher Scientific) using the Invitrogen QubitTM 1X dsDNA High Sensitivity Assay Kit (Thermo Fisher Scientific, Q33231, Waltham, MA, USA). The library concentrations in ng/µL were converted to 4 nM. The 4 nM library concentrations were normalised to a 2 nM concentration and pooled together into a 1 mL microcentrifuge tube. The 2 nM concentration library pool was further diluted to a final loading concentration of 1000 pM (1 nM) by combining 12 µL of the pooled DNA libraries and 12 µL of Resuspension Buffer (RSB) with Tween 20 (Illumina, 20050639). The Illumina 10 nM 5% PhiX Control v3 (Illumina, 15017666) was also diluted to 1000 pM by combining 2 µL of PhiX and 18 µL of RSB with Tween 20. A final volume of 24 µL of the library pool was spiked with 1 µL of 1 nM PhiX control v3. A final sequencing volume of 20 µL was loaded into the Illumina® NextSeq 2000 Sequencing platform (Illumina, San Diego, CA, USA) where there was an onboard denaturation of the spiked library pool and sequenced for 600 cycles using the Illumina NextSeq^TM^ 1000/2000 XLEAP-SBS^TM^ P1 Reagent Cartridge (Illumina, 20101841) to generate 150bp x 2 paired-end reads. This was done at the Next-Generation Sequencing Unit of the University of the Free State, Bloemfontein, South Africa.

### 2.8. Viral metagenomic data analysis

The raw demultiplexed FASTQ reads obtained from the Illumina platform were uploaded into the Chan Zuckerberg ID (CZ ID), a cloud-based, open-source bioinformatics platform pipeline (https://czid.org) (Kalantar et al., 2020) designed for metagenomic analysis. It is also used to detect and track infectious diseases and identify virus types in biological samples. The CD IZ mNGS pipeline preprocesses the uploaded reads and then proceeds to assembly-based alignment to generate taxonomic relative abundance profiles for each sample (Lu et al., 2024). Briefly, the raw FASTQ reads underwent quality control and host filtering. The low-quality and low complexity reads were first removed with Fastp (Chen et al., 2018), host reads were removed with Bowtie2 (Langmead and Salzberg , 2012) and HISAT2 (Kim et al., 2019). The duplicate reads were filtered using CZID-dedup (https://github.com/chnazuckerberg/czid-dedup). The non-host reads output by the quality- and host-filtering steps were aligned to the NCBI nucleotide (NT) and protein (NR) databases using Minimap2 (Li, 2018) and DIAMOND (Buchfink et al., 2015), respectively, to identify possible short-read alignments. Reads were then assembled into contigs using SPADES (Bankevich et al., 2012) and contigs were realigned to the set of possible accessions using Basic Local Alignment Search Tool (BLAST) to improve accuracy. Finally, the alignments were used to identify taxa of origin, which are tallied into relative abundance estimates (Kalantar et al., 2020).

### 2.9. Statistical analysis

The virome diversity and abundance across the study groups were analysed at the species level using both alpha and beta metrics. Participants were categorised into three cohorts: malnourished children with gastroenteritis, non-malnourished children with gastroenteritis and lastly children without gastroenteritis and malnutrition. To minimise biases resulting from uneven sequencing depth across libraries, viral reads were normalised to reads per million (RPM) by dividing the number of reads assigned to each viral species by the total number of reads within the corresponding library and multiplying by one million. Alpha diversity, representing within-sample diversity, was assessed using observed richness (Sobs), Shannon, and Simpson indices based on the viral species composition of each sample. Beta diversity, representing between-sample diversity, was evaluated using the principal coordinate analysis (PCoA) generated from the Bray-Curtis dissimilarity index (Bray and Curtis, 1957) and Jaccard presence-absence distance matrices (Verma and Aggarwal, 2020). Alpha diversity among study groups were analysed using the non-parametric Kruskal–Wallis test. The Bray-Curtis dissimilarity matrices were used to evaluate beta diversity from the viral abundance of each sample. Differences in viral community composition across groups were assessed using permutational multivariate analysis of variance (PERMANOVA) using the adonis test with 1000 permutations.

### 2.10. Data visualisation

The data visualisations and subsequent statistical analyses were performed on R software (version 4.5.1). Viral read abundance and taxonomic tables used as input files were formatted, analysed, and visualized using the R packages vegan (Oksanen et al., 2015), ggplot2 (Wickham et al., 2016), cowplot (Wilke et al., 2019), pheatmap (Kolde and Kolde, 2018), and tidyverse (Wickham et al., 2019).

## 3.0. Results

### 3.1. An overview of the study population

A total of 69 samples were processed using the metagenomic sequencing-approach. Overall, 38 (55.1%) and 31 (44.9%) samples were from female and male participants, respectively. The study comprised three cohorts of 23 samples each after initial exclusions. The malnourished children with gastroenteritis cohort included 11 (47.8%) females and 12 (52.2%) males, with a mean age of 12.1 months, a weight of 6.0 kg and a height of 66.1 cm. The non-malnourished children with gastroenteritis cohort comprised 16 (69.6%) female participants and 7 (30.4%) male participants, with a mean age, weight, and height of 13.0 months, 7.1 kg, and 67.3 cm, respectively. The third cohort, consisting of children without gastroenteritis and malnutrition (healthy controls), included 11 (47.8%) females, 12 (52.2%) males, with a mean age, weight, and height of 13.7 months, 8.6 kg and 69.5 cm, respectively (Supplementary Table S2).

### 3.2. Overview of metagenomic data

Of the 69 samples analysed, sequencing failed for 2 samples and one additional sample yielded no reads assignable to viruses. Consequently, 66 samples were retained for downstream virome analysis. Metagenomic sequencing of 67 samples generated a total of 138,701,122 paired-end reads (151 bp per read) using the Illumina NextSeq 2000 platform, with an average of 2,070,166 raw reads generated per sample and 1,525,284 reads per sample passing quality filters. Following bioinformatic analysis, 83,732,042 (60.4%) of the total raw reads were classified, while 54,969,080 (39.6%) remained unclassified. Of the classified reads, 76,185,944 (91.0%) were assigned to bacteria, 5,375,381 (6.4%) to viruses, and 2,170,717 (2.6%) to other microorganisms (Table 1).

**Table 1.** Metagenomic overview of sequenced reads.

| **Category** | **Total number of raw reads** | **Average reads per sample** | **Average reads passing filters per sample** | **Classified reads** | **Unclassified reads** | **Bacterial reads** | **Viral reads** | **Others (i.e., host, etc)** |
|---|---|---|---|---|---|---|---|---|
| **Reads Count** | **138,701,122** | 2,070,166 | 1,525,284 | 83,732,042 | 54,969,080 | 76,185,944 | 5,375,381 | 2,170,717 |
| **Percentage/Total raw reads** |  |  |  | 60.4% | 39.6% | 54.9% | 3.9% | 1.6% |
| **Percentage/Total classified reads** |  |  |  |  |  | 91.0% | 6.4% | 2.6% |

### 3.3. Distribution of the number of viral species reads based on established natural host specificity

Among the viral reads, 5,046,194 (93.9%) were assigned to eukaryotic viruses and 329,187 (6.1%) to bacteriophages. Eukaryotic viral reads were further classified into natural host type based on the viral species. Viral species from human host type were the most abundant: 4,886,045 (96.8%), followed by those from non-human mammal host:153,572 (3.04%), plant host: 3,275 (0.065%), protist host: 2,918 (0.058%), fungal host: 296 (0.006%), avian host: 91 (0.002%), arthropod host: 24 (0.0005%) and unknown host: 17 (0.0003%) respectively **(Fig. S1).**

### 3.4. Detected Eukaryotic viral families

Analysis of 66 human faecal samples yielded 5,046,194 eukaryotic viral reads, which were classified into 39 viral families, 71 viral genera, 243 viral species and some unclassified viruses. The viral families were further grouped into 9 DNA viral families *(Adenoviridae, Herpesviridae, Papillomaviridae, Polyomaviridae, Mimiviridae, Annelloviridae, Parvoviridae* and *Circoviridae*) and 30 RNA viral families (*Astroviridae, Caliciviridae, Picornaviridae, Coronaviridae, Matonaviridae, Alphaflexivirdae, Betaflexiviridae, Bromoviridae, Closteroviridae, Luteoviridae, Potyviridae, Solemoviridae, Tombusviridae, Virgaviridae, Botourmiaviridae, Mitoviridae, Narnaviridae, Dicistroviridae, Paramyxoviridae, Pneumoviridae, Sedoreoviridae, Amalgaviridae, Botybirnaviridae, Chrysoviridae, Endornaviridae, Partitiviridae, Totiviridae, Picobirnaviridae, Reoviridae and Retroviridae*), detection frequency per group, overall detection in all 66 samples, and natural host type respectively **(Tables S4, S5, and S6).**

#### 3.4.1. Frequency distribution of detected viral families

The prevalence of both enteric and non-enteric viral families was investigated in all 66 samples. Among the human enteric families, *Sedoreoviridae* was the most prevalent, detected in 59/66 (89.4%) samples. This was followed by *Picornaviridae* detected in 56/66 (84.9%), *Caliciviridae* in 35/66 (53.0%), *Astroviridae* in 33/66 (50.0%), and, lastly, *Adenoviridae* in 17/66 (25.6%) **(Fig. 1).**

**Fig. 1.**
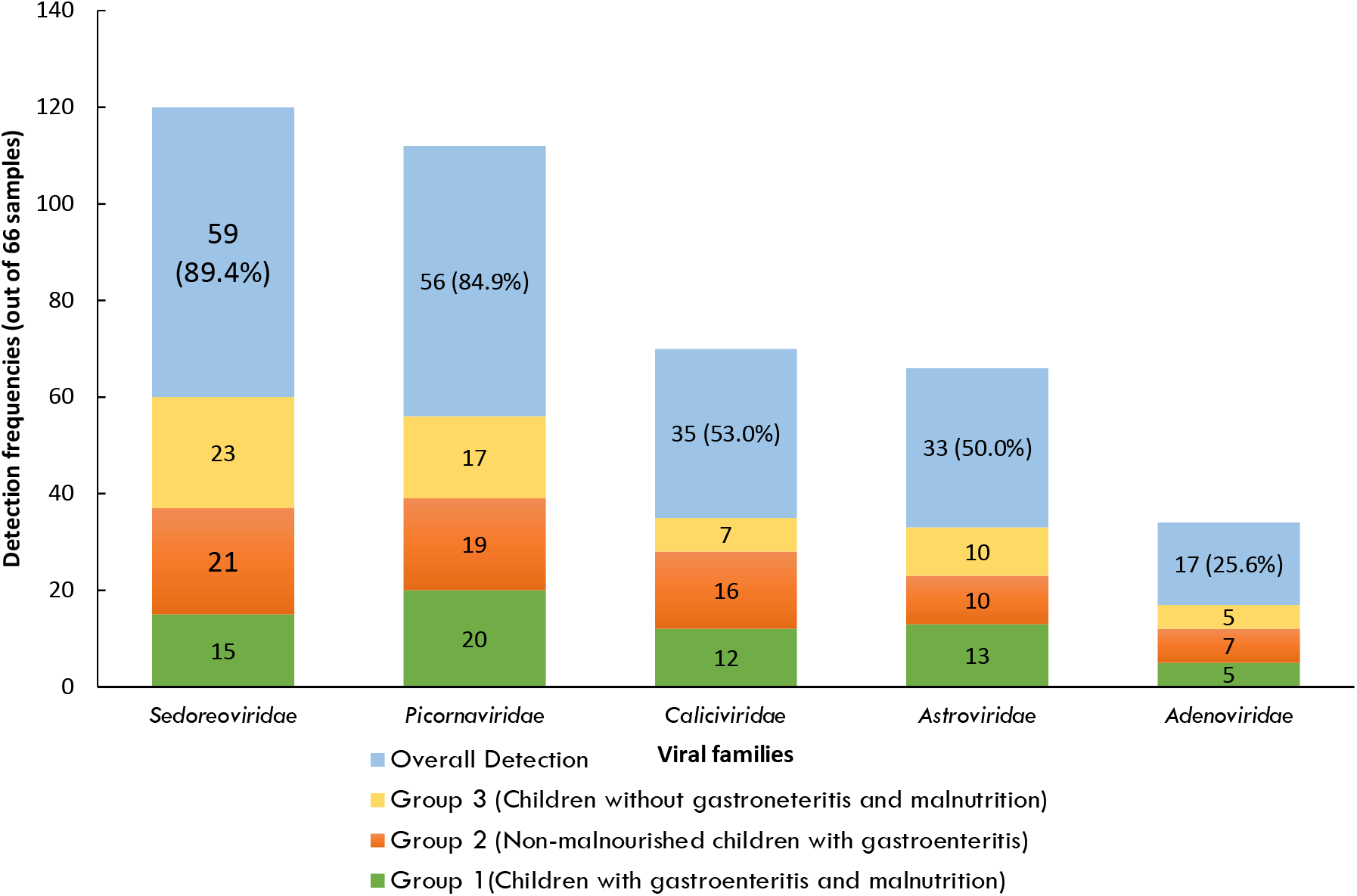
Detection frequencies of eukaryotic enteric viral families across the three study groups: Group 1(Children with gastroenteritis and malnutrition), Group 2 (Non-malnourished children with gastroenteritis) and Group 3 (Children without gastroenteritis and malnutrition).

Viral families not directly associated with gastroenteritis in human were also detected in the infants’ gut at different detection frequencies, with a predominance of *Retroviridae* 39/66 (59.1%), *Picobirnaviridae* 31/66 (46.9%), *Parvoviridae* 25/66 (37.9%), *Virgaviridae* 24/66 (36.4), and *Anelloviridae* 21/66 (31.8%) **(Fig. S2).**

### 3.5. Comparative eukaryotic virome composition and relative abundance across the three study cohorts

#### 3.5.1. Malnourished children with gastroenteritis

A total of 1,536,098 eukaryotic viral reads were generated from 21 samples retained for downstream analysis, accounting for 30.4% of the total eukaryotic reads. A total of 19 classified viral families, 33 genera and 126 viral species were identified in the cohort comprised of malnourished children with gastroenteritis. Based on genome organisation, two viral families had double-stranded DNA (dsDNA) genomes: *Adenoviridae* and *Herpesviridae*. *Anelloviridae* was the only detected family with a circular negative-sense single-stranded DNA (-ssDNA) genome, while *Parvoviridae* was the lone viral family with a single-stranded DNA (ssDNA) genome. Among RNA viruses, eight viral families possess positive-sense single-stranded RNA (+ssRNA) genome, including *Astroviridae, Bromoviridae, Caliciviridae, Coronaviridae, Narnaviridae, Picornaviridae, Solemoviridae,* and *Virgaviridae.* Furthermore, five viral families with a double-stranded RNA (dsRNA) genome were detected, including *Chrysoviridae, Partitiviridae, Picobirnaviridae, Sedoreoviridae,* and *Totiviridae. Paramyxoviridae* was the only detected family with a negative-sense single-stranded RNA (-ssRNA), while *Retroviridae* was the only family identified with a positive-sense single-stranded RNA genome with reverse transcription (+ssRNA-RT) **(Table S4).**

*Caliciviridae* was the most abundant enteric viral family with 1,180,936 reads accounting for 76.88% of the total eukaryotic viral reads for this group and was detected in 12/21 samples. This was followed by *Picornaviridae* with 280,579 reads (18.27%), detected in 20/21 samples, *Sedoreoviridae* with 31,164 reads (2.03%), and detected in 15/21 samples, *Astroviridae* with 21,421 reads (1.29%), detected in 12/21 samples and lastly *Adenoviridae* family with 1,251 reads (0.08%) detected in 5/21 samples **(Fig. 2.; Table S4).** Other viral families not directly linked with gastroenteritis were also detected at different frequencies and presented in order of most read abundance, *Picobirnaviridae, Parvoviridae, Coronaviridae, Virgaviridae, Retroviridae, Anelloviridae, Totiviridae, Herpesviridae, Paramyxoviridae, Partitiviridae, Solemoviridae, Narnaviridae, Chrysoviridae* and *Bromoviridae* **(Fig. 2.; Table S4).**

**Fig. 2.**
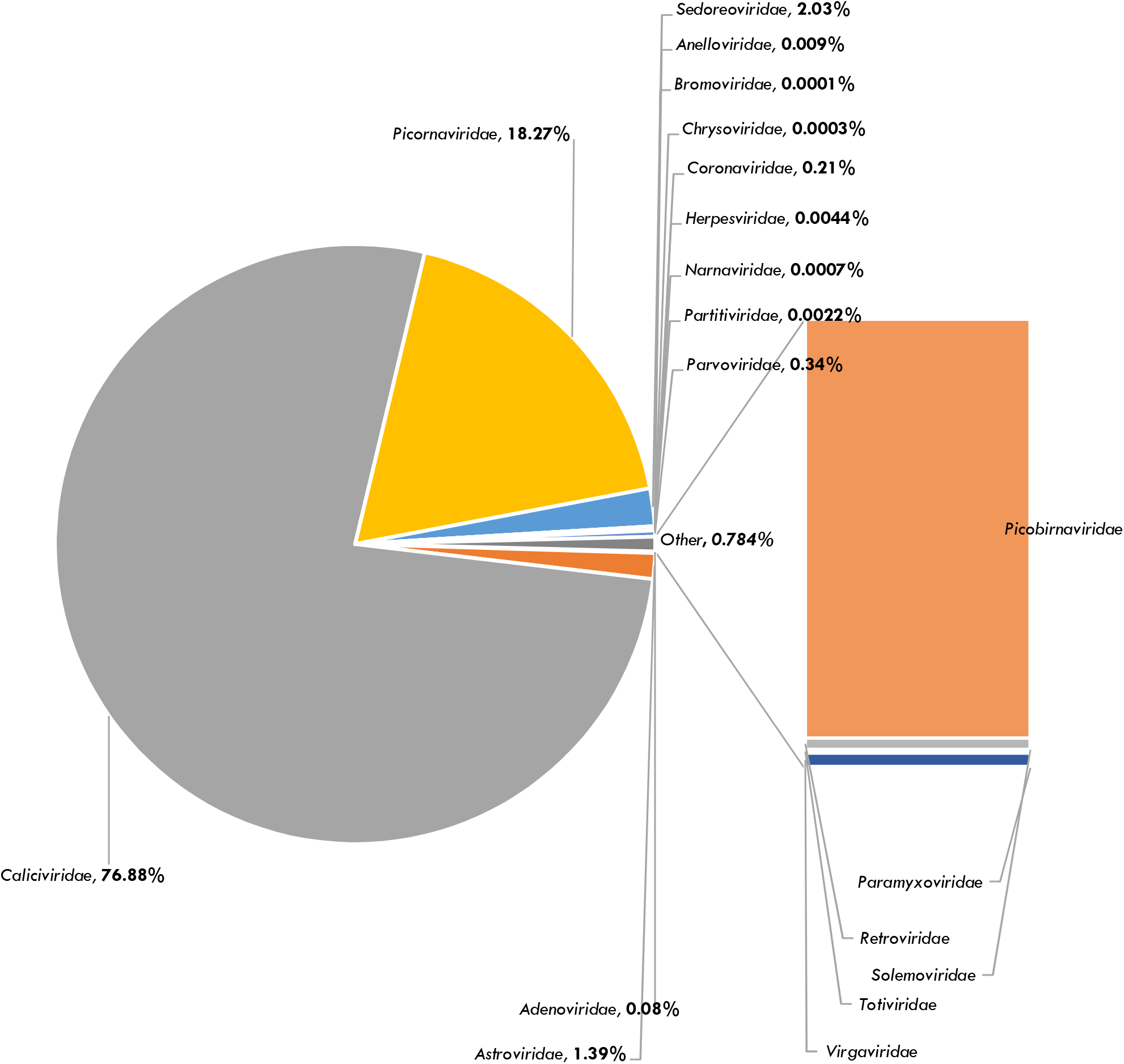
Taxonomic distribution of eukaryotic viral families identified by metagenomic sequencing in malnourished children with gastroenteritis.

At the genus level, the eukaryotic viral reads were classified into 33 viral genera. Ten of these viral genera directly associated with gastroenteritis in children accounted for 98.6% of the reads, with a predominance of *Sapovirus,* a member of the *Caliciviridae* family, with 1,180,715 assigned reads constituting 76.86% of the reads. Also abundant was the genus *Enterovirus* within the *Picornaviridae* family, with 240,727 (15.6%). Among the enteric genera within the *Picornaviridae* family, the detected genera were *Salivirus* (36,377 reads), *Parechovirus* (3,055 reads), *Cosavirus* (284 reads), and *Kobuvirus* (1 read; **Table S4).** Other well-known enteric genera circulating among this cohort included *Rotavirus* with 31,164 reads representing only 2.03% of the reads, *Mamastrovirus* with 21,421 reads 1.39%, *Mastadenovirus* with 1,251 (0.08%) and *Norovirus* with 221 reads (0.01%). Additionally, *Alphatorquevirus*, *Betatorquevirus*, *Gammatorquevirus, Betacoronavirus, Bocaparvovirus, Cytomegalovirus, Dependoparvovirus, Lentivirus, Orthopicobirnavirus, Picobirnavirus, Respirovirus, Roseolovirus, Sapelovirus, and Torovirus, all mammalian-infecting viral genera with no direct relationship with gastrointestinal tract infections, were detected at different read abundances*. *Alphachrysovirus*, *Gammapartitivirus, Totivirus* and *Victorivirus* all from fungi host and *Cryspovirus* of protists host were identified. Plants infecting genera *Ilarvirus* from the *Bromoviridae* family, *Sobemovirus* from the *Solemoviridae* family, and *Tobamovirus* belonging to the *Virgaviridae* family were the most abundant, with 371 reads present **(Table S4).** Some detected viral sequences could not be assigned to specific genera.

Viral species in this group associated with gastroenteritis in children below the age of five included *Sapovirus, Adenoviruses, Astroviruses, Norovirus, Cosaviruses A* and *D, Enteroviruses A, B,* and *C, Kobuvirus, Parechovirus A, Salivirus A, Rotavirus A, Mastadenoviruses F,* and *Mamastrovirus 1.* Among them, *Sapovirus* was the most abundant, with 1,180,715 reads, over a million in one sample, and detected in 8/21 samples. This was followed by *Enterovirus B, Rotavirus A, Salivirus A, Human enterovirus, Enterovirus C, Enterovirus A, Parechovirus A, Norovirus and Enterovirus C*. Other human enteric viral species were also identified **(Table S4).** Human rhinovirus sp., Rhinovirus A, B, and C, Betacoronavirus 1, viral species responsible for respiratory infections in humans, were present. Interestingly, viral species primarily known to infect mammals, plants, fungi and protist hosts were also detected in faecal samples collected **(Table S4).**

#### 3.5.2. Non-malnourished children with gastroenteritis

For the second study group, consisting of non-malnourished children with gastroenteritis, a total of 1,879,303 reads were generated from 22 faecal samples, representing 37.24% of the total eukaryotic viral reads. A total of 32 classified viral families, comprising 52 genera and 144 viral species, were identified. Based on the genome organisation, 2 viral families possessed a double-stranded DNA (dsDNA) genome, and these included *Adenoviridae* and *Polyomaviridae*, 1 family with a circular negative-sense single-stranded DNA (-ssDNA) genome structure; *Anelloviridae,* 2 viral families with a single-stranded DNA (ssDNA) genome structure; *Parvoviridae,* and *Circoviridae.* Among RNA viruses, 16 families possessed a positive-sense single-stranded RNA (+ssRNA) genome, including *Alphaflexiviridae, Astroviridae, Betaflexviridae, Botourmiaviridae, Caliciviridae, Closteroviridae, Coronaviridae, Dicistroviridae, Luteoviridae, Matonaviridae, Narnaviridae, Picornaviridae, Potyviridae, Solemoviridae,* Tombusviridae, and *Virgaviridae.* Additionally, 9 Viral families with a double-stranded RNA (dsRNA) genome were detected, including *Amalgaviridae*, *Chrysoviridae, Botybirnaviridae, Endornaviridae, Partitiviridae, Picobirnaviridae, Reoviridae, Sedoreoviridae,* and *Totiviridae. Pneumoviridae* was the only family identified with *a* negative-sense single-stranded RNA (-ssRNA) genome, while *Retroviridae* was the only detected viral family possessing a positive-sense single-stranded RNA genome with reverse transcription (+ssRNA-RT) **(Table S5).**

In contrast with the malnourished children with gastroenteritis group, *Sedoreoviridae* predominated the enteric viral families in this cohort, with 1,412,548 reads, representing 75.16% of total eukaryotic viral reads. *Picornaviridae* was the next most abundant family with 296,899 reads (15.79%), followed by *Astroviridae*, with 47,339 reads (2.52%), *Adenoviridae* with 7,642 reads (0.41%), and *Caliciviridae with* 1,926 reads (0.10%) **(Fig. 3; Table S5).** Twenty-seven viral families not commonly implicated in gastroenteritis were also detected, including *Anelloviridae, Coronaviridae, Matonaviridae, Parvoviridae*, *Picobirnaviridae, Pneumoviridae, Polyomaviridae, Reoviridae,* and *Retroviridae,* all mammalian-infecting viruses. Plant-infecting viral families included *Alphaflexiviridae, Amalgaviridae, Betaflexiviridae, Closteroviridae, Endornaviridae, Luteoviridae, Potyviridae, Solemoviridae, Tombusviridae,* and *Virgaviridae.* Fungal-infecting viral families comprised *Chrysoviridae, Botourmiaviridae, Botybirnaviridae, Narnaviridae, Partitiviridae* and *Totiviridae*. *Circoviridae* was the only avian-infecting family detected, while *Dicistroviridae* was identified as the only insect-infecting viral family **(Fig. 3; Table S5).**

**Fig. 3.**
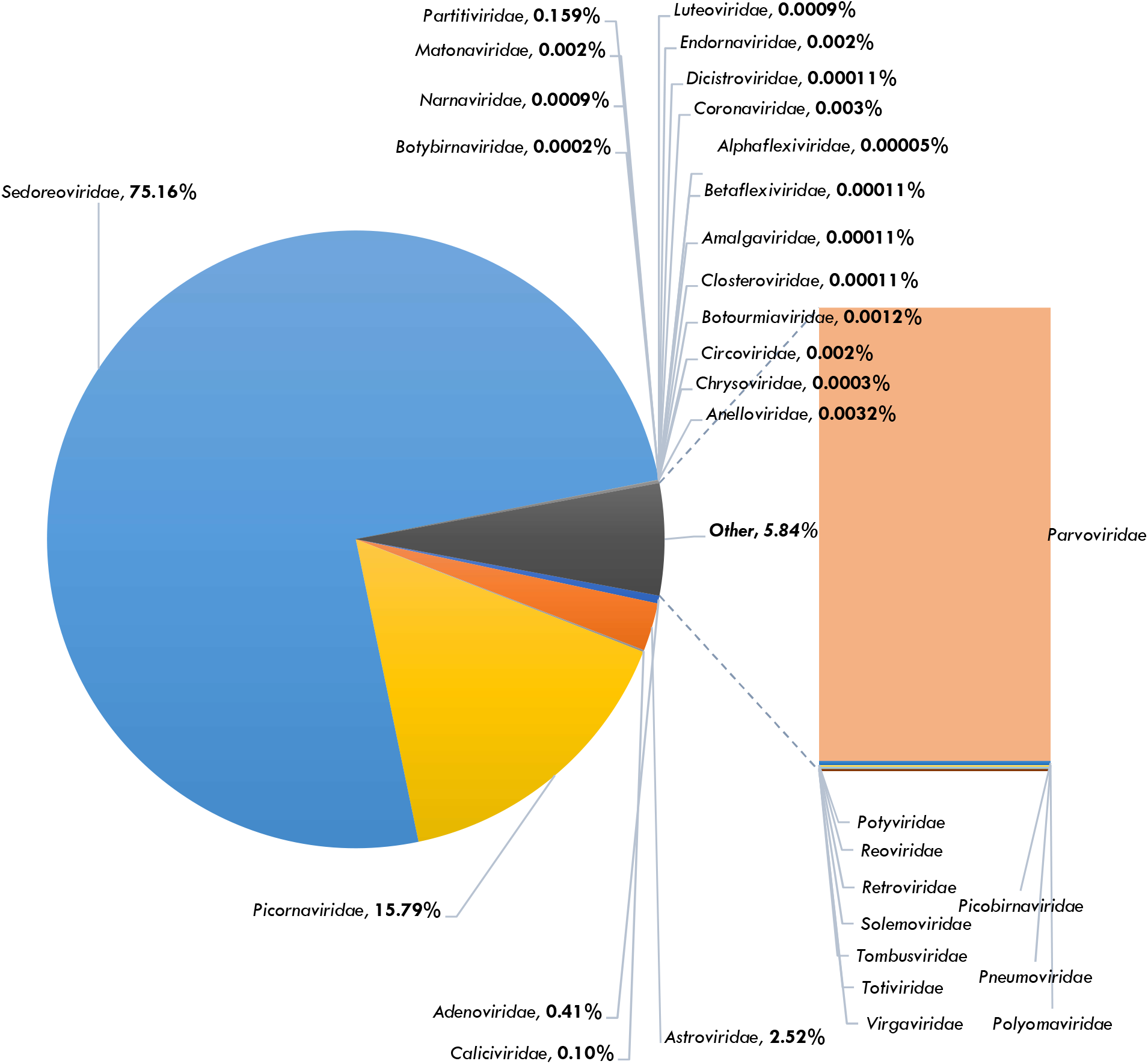
Taxonomic distribution of eukaryotic viral families identified from sequenced reads in non-malnourished children with gastroenteritis.

Among the 52 classified eukaryotic viral genera in this cohort, nine were enteric, with *Rotavirus* representing the largest proportion of the viral reads (1,412,548; 75.16%) and detected in 21/22 samples. Other commonly detected human enteric viral genera included *Mamastrovirus, Mastadenovirus, Sapovirus*, and Norovirus, which were present at varying abundances, representing 3.03% of the reads. Interestingly, *Sapovirus* was detected at a lower relative abundance observed in malnourished children with gastroenteritis. *Cardiovirus, Cosavirus, Enterovirus* and *Parechovirus*, all enteric genera within the *Picornaviridae* family were also identified, with *Cardiovirus* being the most abundant with 143,150 reads. Non-enteric mammalian-infecting viral genera detected at different reads abundance included, *Alphatorquevirus*, *Betapolyomavirus*, *Betaretrovirus*, *Betatorquevirus*, *Gammaretrovirus*, *Gammatorquevirus Alphacoronavirus, Betacoronavirus*, *Bocaparvovirus, Copiparvovirus*, *Dependoparvovirus*, *Hepatovirus*, *Lentivirus, Metapneumovirus*, *Orthoreovirus*, *Orthopicobirnavirus, Picobirnavirus, Roseolovirus*, *Rubivirus, Sapelovirus, and Torovirus*. *Aveparvovirus* was the only Avian-infecting viral genus identified in this group. A large number of plant-infecting and fungal-infecting genera were also identified, although some detected viral sequences remained unclassified at the genus level **(Table S4).**

At the species level, a diverse range of enteric viral species belonging to the Masta*denovirus, Mamastrovirus, Cardiovirus, Cosavirus, Enterovirus, Parechovirus* and *Rotavirus* genera were detected at varying read counts, with *Rotavirus A* emerging as the predominant species, accounting for 1,406,776 reads and detected in 21/22 samples. This was followed by *Cardiovirus B*, *Enterovirus B, Mamastrovirus 1, Parechovirus A, Enteroviruses C, Cardiovirus D, Human mastadenovirus,* and *Human astrovirus. Sapovirus, Norovirus, Human mastadenoviruses B and D, Rotavirus C, Human enterovirus, Enterovirus A, and Mamastrovirus 3* were detected at comparatively lower read abundances of 1,796, 130, 16, 24, 14, 844, 61, 15, and 4, respectively **(Table S5).** Human respiratory viral species, including *severe acute respiratory syndrome-related coronavirus*, *Human rhinovirus species, and Rhinovirus A, B,* and *C,* were also detected. In addition, *Hepatovirus A*, a viral species within the *Picornaviridae* family, *Metapneumovirus hominis* within the *Pneumoviridae* family, *and Betapolyomavirus hominis within the Polyomaviridae family, all known human-infecting pathogens not associated with gastroenteritis,* were identified. Additionally, viral species of animal, plant, fungal and protist hosts’ specificity were detected in this study group **(Table S5).**

#### 3.5.3. Children without gastroenteritis and malnutrition

A total of 1,630,793 eukaryotic viral reads were generated from 23 samples, representing 32.32% of the total eukaryotic reads, and were classified into 23 viral families, 32 genera, and 83 viral species. Based on genome organisation, five viral families possessed a double-stranded DNA (dsDNA) genome, namely *Adenoviridae, Herpesviridae, Mimiviridae, Orthoherpesviridae, and Papillomaviridae. Anelloviridae* was the only detected viral family with a circular negative-sense single-stranded DNA (-ssDNA) genome, while *Parvoviridae* the only family with a single-stranded DNA (ssDNA) genome. Among RNA viruses, 10 viral families possessed a positive-sense single-stranded RNA (+ssRNA) genome, including *Astroviridae, Botourmiaviridae, Caliciviridae, Dicistroviridae, Mitoviridae, Narnaviridae, Picornaviridae, Potyviridae, Solemoviridae,* and *Virgaviridae.* Five viral families with a double-stranded RNA (dsRNA) genome were identified, including *Endornaviridae, Partitiviridae, Picobirnaviridae, Sedoreoviridae,* and *Totiviridae,* while *Retroviridae* was the only detected viral family possessing a positive-sense single-stranded RNA genome with reverse transcription (+ssRNA-RT) **(Table S6).**

Similar to the non-malnourished children with gastroenteritis cohort, *Sedoreoviridae* was the most abundant enteric viral family in this group, with 1,566,489 reads, representing 96.06% of the eukaryotic viral reads. More enteric viral families identified included *Picornaviridae* with 60,899 reads (3.71%), *Astroviridae*, with 265 reads (0.02%), *Caliciviridae* with 104 reads (0.01%), and *Adenoviridae* with 50 reads (0.003%) **(Fig.4; Table S6).** Viral families not commonly implicated in gastroenteritis were also detected, including *Anelloviridae, Herpesviridae, Orthoherpesviridae, Papillomaviridae, Parvoviridae*, *Picobirnaviridae,* and *Retroviridae,* all of which are mammalian-infecting viruses. Plant-infecting viral families included *Endornaviridae, Potyviridae, Solemoviridae,* and *Virgaviridae.* Fungal-infecting viral families comprised *Botourmiaviridae, Narnaviridae, Partitiviridae* and *Totiviridae*, while *Dicistroviridae* was the only insect-infecting family detected, and *Mimiviridae* the only protist-infecting family identified **(Fig. 4; Table S6).**

**Fig. 4.**
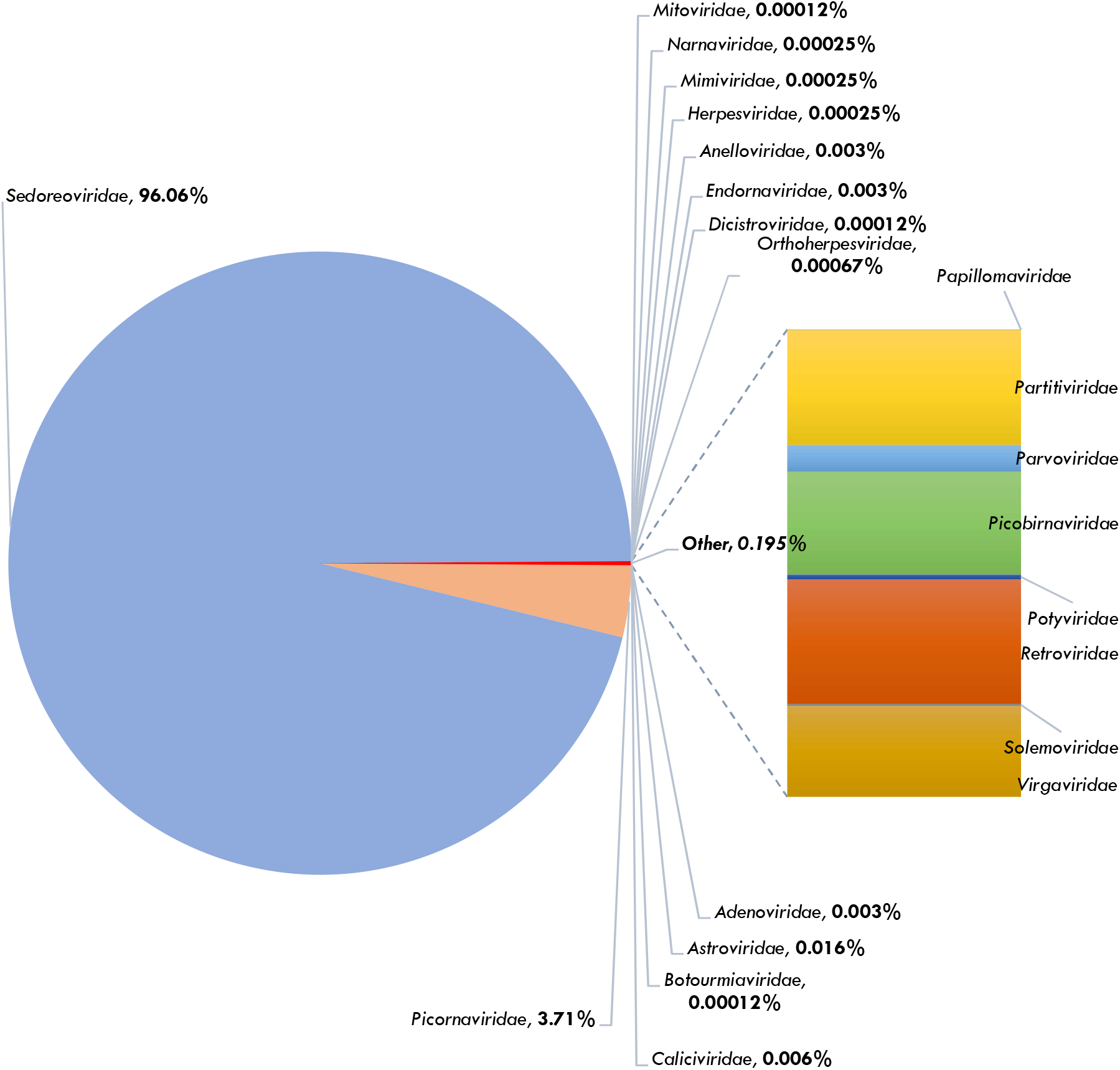
Taxonomic profiling of eukaryotic viral families related sequencing reads in children without gastroenteritis and malnutrition.

Among the 32 classified eukaryotic viral genera in this cohort, seven were enteric, with *Rotavirus,* the lone genus within the *Sedoreoviridae* family representing the largest proportion of the viral reads, 1,566,489 (96.06%), and was detected in all 23 samples. Other commonly detected human enteric viral genera in decreasing order of read abundance included *Enterovirus* with 59,991 reads (3.68%), *Parechovirus* with 472 reads (0.03%), *Mamastrovirus* with 265 reads (0.02%), *Norovirus* with 92 reads (0.005%), *Mastadenovirus* with 50 reads (0.003%) and *Sapovirus* with 12 reads (0.0007%). Non-enteric mammalian-infecting viral genera detected at different read abundance included, *Alphatorquevirus*, *Cytomegalovirus, Lymphocryptovirus, Roseolovirus*, *Muromegalovirus, Alphapapillomavirus,* Chaphamaparvovirus, *Dependoparvovirus*, *Gammaretrovirus, Lentivirus, Orthopicobirnavirus, Picobirnavirus,* and *Sapelovirus.* Plant-infecting viral genera identified included *Alphaendornavirus, Polerovirus, Potyvirus, Sobemovirus* and *Tobamovirus,* while *Ourmiavirus, Totivirus,* and *Victorivirus* were the fungal-infecting viral genera protected. Protist-infecting genera *included Megavirus, Mimivirus* and *Theiavirus, whereas Gyrovirus* was the only avian-infecting viral genus detected. Some detected viral sequences remained unclassified at the genus level **(Table S6).**

At the species level, metagenomic analysis identified 83 eukaryotic viral species in this study cohort. A wide range of enteric viral species within the Masta*denovirus, Mamastrovirus, Enterovirus, Parechovirus* and *Rotavirus* genera were detected at varying read counts, with *Rotavirus A* predominating, accounting for 1,565,857 reads and detected in all 23 samples; notably, over one million reads were detected in a single sample This was followed by *Enterovirus C, Parechovirus A, Human enterovirus*, *Norovirus, Mamastrovirus 1, Human mastadenovirus F, Human mastadenovirus B, Sapovirus, and Human mastadenovirus A* **(Table S6).** Non-enteric human viral species identified within the *Herpesviridae* family included *Cytomegalovirus humanbeta5, Lympocryptovirus humangamma4* and *Human betaherpesvirus 6*. Human respiratory viral species, including *Antwerp rhinovirus 98/99*, *Human rhinovirus sp., Rhinovirus A* and *C* were also detected. Additionally, viral species with animal, plant, fungal, avian and protist host specificity were identified in this study group **(Table S6).**

#### 3.5.4. Species diversity and abundance of human enteric viruses across study cohorts

Analysis of the metagenomic data from all 66 faecal samples yielded a total of 5,046,194 eukaryotic viral reads. Human viral reads predominated in faeces with 4,886,045 reads accounting for 96.83% of the total eukaryotic viral reads. These reads were further classified into 243 viral species, with mammalian viruses representing the predominant group. Among the human-infecting enteric viruses, *Rotavirus A* was the most abundant across all study cohorts, with 3,003,785 reads accounting for 59.5% of the total eukaryotic reads and a detection frequency of 59/66 (89.4%). *Sapovirus* was the second most abundant enteric virus with of a total of 1,182,523 reads, representing 23.4% of the total eukaryotic viral read, and was detected in 20/66 samples (30.3%). Other human enteric viral species were also detected at varying read abundances and frequencies across the study cohorts **(Fig. S2).**

#### 3.5.5. Variations in the faecal virome composition, abundance and diversity across study cohorts

Based on relative standardised abundance, most sequence reads across all libraries were assigned to RNA viruses, with their relative abundance of the 20 most abundant viral species across study the groups presented in **Fig. 5**. These results reveal distinct differences in virome composition among the three cohorts accompanied by inter-sample variability.

**Fig. 5.**
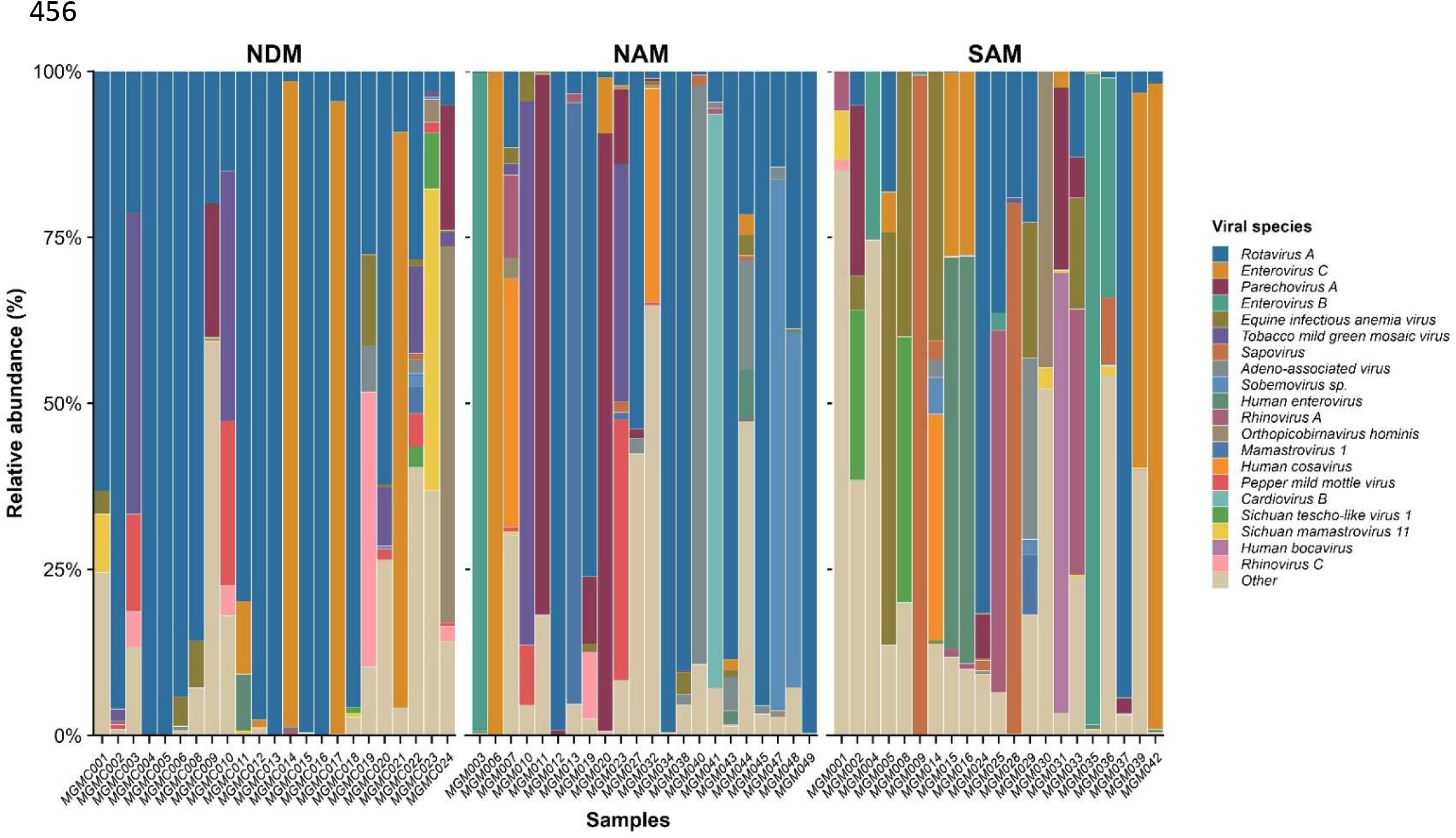
Comparative relative abundance of the 20 most abundant viral species across individual faecal samples from children without gastroenteritis and malnutrition (NDM), non-malnourished children with gastroenteritis (NAM), and malnourished children with gastroenteritis (SAM).

Across the cohorts consisting of children without gastroenteritis and malnutrition, and non-malnourished children with gastroenteritis. Rotavirus A was the predominant viral species in most samples, frequently accounting for a large proportion of the total viral reads. Several samples were almost exclusively dominated by Rotavirus A, whereas others exhibited mixed viral populations comprising *Enterovirus C, Parechovirus A, Rhinovirus A*, *Human bocavirus, Mastadenovirus,* and *Mamastrovirus 1* at varying relative abundances **(Fig. 5).**

In contrast, the malnourished children with gastroenteritis cohort exhibited a more heterogeneous viral composition, with fewer samples dominated by *Rotavirus A*. Instead, several samples showed predominance of *Sapovirus, Enterovirus C, Orthopicobirnavirus hominis*, and *Human bocavirus*, while others contained more evenly distributed viral communities composed of multiple viral species **(Fig. 5).** Animal-associated viruses, including *Equine infectious anaemia virus*, together with plant-associated viruses such as *Tobacco mild green mosaic virus*, were also detected in individual samples, although generally at lower relative abundances.

Still at the species level, *Rotavirus A*, emerged as the most abundant and widespread viral species across nearly all samples and study cohorts. Other human-infecting enteric viruses detected at varying abundances across the cohorts included *Norovirus, Human enterovirus, Enterovirus A, Enterovirus B, Enterovirus C, Enterovirus D, Parechovirus A, Cardiovirus B, Cardiovirus D, Cosavirus A, Cosavirus D, Human astrovirus, Human mastadenovirus A,* and *Human mastadenovirus F* **(Fig. S4).** The non-enteric human viruses included; *Enterovirus D*, *Human astrovirus MLB2, Human mink ovine-like (HMO) astrovirus, Human mastadenovirus B, Human mastadenovirus and Human mastadenovirus D.* Several human respiratory-associated viruses were also detected, albeit at lower relative abundance. These included *Rhinovirus A, Rhinovirus B, Rhinovirus C, Human betacoronavirus, severe acute respiratory syndrome-related coronavirus,* and *Respirovirus laryngotracheitidis.* Several DNA viruses known to infect humans were detected sporadically across samples with variable abundance including *Human bocavirus, Bocaparvovirus primate 1, Betapolyomavirus hominis, TTV-like mini virus, Torque teno virus, Human betaherpesvirus 6* and *Cytomegalovirus humanbeta5* **(Fig. S4).** Notably, animal-associated viruses were also identified, including *Porcine picobirnavirus* and *Equine infectious anaemia virus*. Similarly, several plant-associated viruses were detected, including *Tobacco mild green mosaic virus, Tomato mosaic virus, Tobacco mosaic virus, Tomato mottle mosaic virus* and *Sobemovirus sp.* **(Fig. S4).**

#### 3.5.6. Reads distribution and detection frequency of clinically important human, animal, plant, avian and fungi genera

Comparative analysis of viral genera across the three study cohorts revealed shared and distinct virome profiles, with considerable variability among individual samples. The infant faecal virome was dominated by a small number of highly abundant viral genera, particularly *Rotavirus* and *Sapovirus* **(Fig. 6).** *Rotavirus* was the most abundant and widely distributed genus, with 3,101,189 reads detected in 59/66 (89.4%) samples, and was the only genus detected in all 23/23 (100%) samples from the NDM cohort. *Sapovirus* was detected across all three cohorts, with 1,182,523 reads and an overall detection frequency of 20/66 (30.3%), with the highest detection observed among malnourished children with gastroenteritis. Other human enteric viral genera, including *Enterovirus*, *Mamastrovirus*, *Parechovirus*, *Mastadenovirus*, and *Norovirus*, were detected across all three cohorts at varying abundances and frequencies. In contrast, *Cardiovirus* was detected exclusively in the non-malnourished gastroenteritis cohort, while *Salivirus* and *Kobuvirus* were detected only in malnourished children with gastroenteritis. *Cosavirus* occurred in both gastroenteritis cohorts **(Fig. 6).** Several non-enteric human associated viral genera were also detected, including *Dependoparvovirus*, *Lentivirus*, *Orthopicobirnavirus*, *Alphatorquevirus*, *Betatorquevirus*, and *Gammatorquevirus*. Respiratory-associated genera, including *Betacoronavirus*, *Respirovirus*, and *Metapneumovirus*, were detected at relatively low abundance and frequency. The virome also contained animal, plant, fungal, and avian associated viral genera, generally at low abundance. Plant-associated genera included *Tobamovirus*, *Sobemovirus*, *Endornavirus*, *Potyvirus*, *Alphaendornavirus*, *Alphapartitivirus*, and *Tombusvirus*, with *Tobamovirus* being the most frequently detected (17/66; 25.8%) and present across all cohorts. Among fungal-associated viruses, *Totivirus* was the most frequently detected (19/66) and occurred across all three cohorts. Animal-associated genera included *Picobirnavirus*, *Sapelovirus*, *Circovirus*, *Muromegalovirus*, and *Orthoreovirus*, while the avian-associated genus *Gyrovirus* was detected less frequently.

**Fig. 6.**
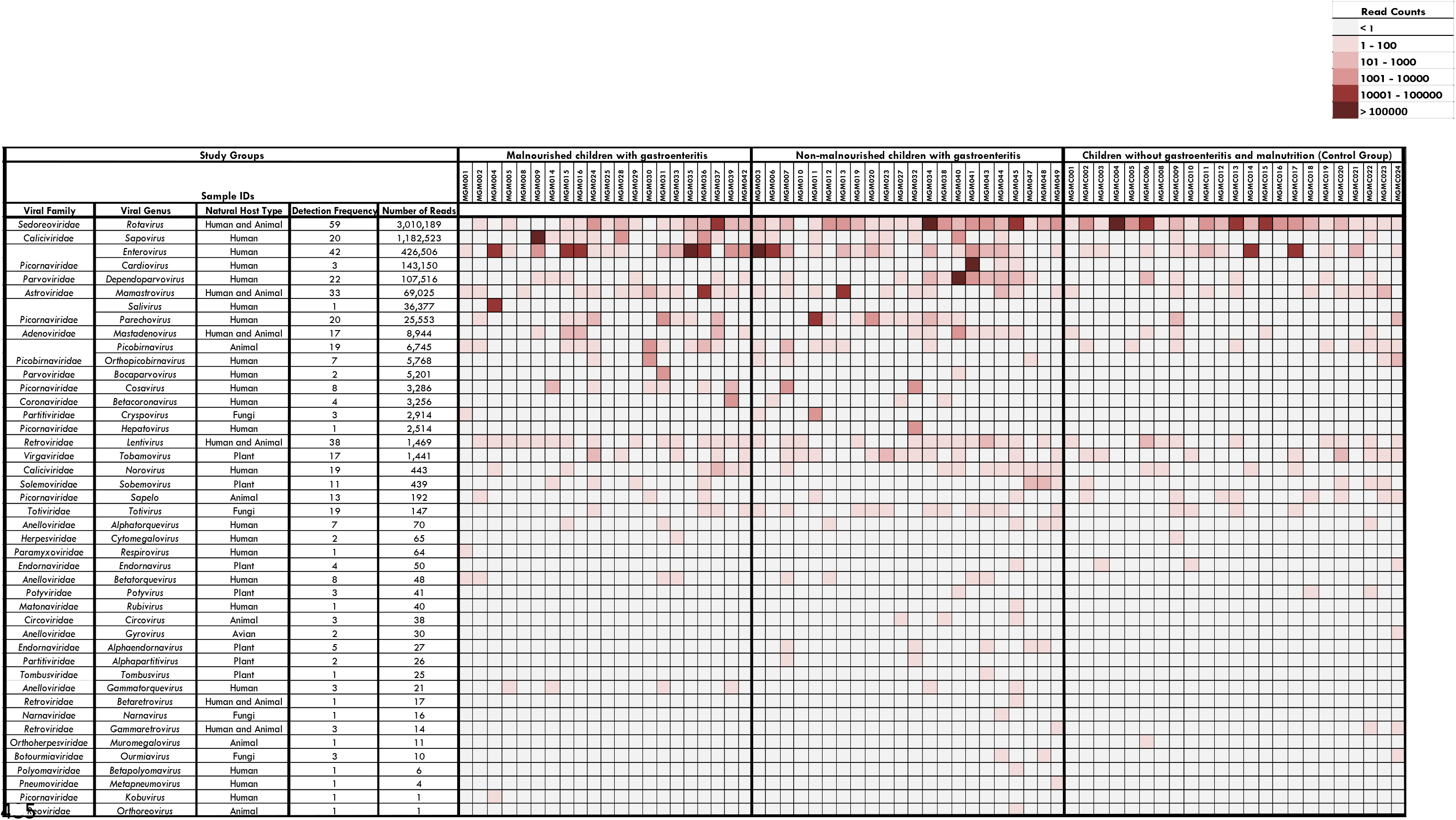
Heatmap illustrating the distribution and abundance of viral genera across individual faecal samples from the three study cohorts: malnourished children with gastroenteritis (SAM), non-malnourished children with gastroenteritis (NAM), and children without gastroenteritis and malnutrition NDM. Viral genera are represented in rows and individual samples in columns, with samples grouped according to study cohort. Colour intensity represents the relative abundance of sequencing reads assigned to each viral genus, with increasing colour intensity indicating higher read abundance.

Overall, the findings demonstrate that the infant faecal virome was predominantly composed of a limited number of highly abundant human enteric viral genera, particularly *Rotavirus* and *Sapovirus*, alongside a diverse range of human non-enteric and non-human-associated viruses occurring at lower abundance and varying frequencies (**Fig. 6**).

#### 3.5.7. Co-occurrences of Common human gastroenteritis-associated viruses

Among the children, at least one enteric viral genus associated with gastroenteritis was detected in stool samples, including *Rotavirus* (RV), *Mastadenovirus (AdV), Sapovirus* (SaV*), Norovirus* (NoV*), Mamastrovirus* (AstV), *Enterovirus* (EV), *and Parechovirus* (PeV) **(Fig. 7)**.

**Fig. 7.**
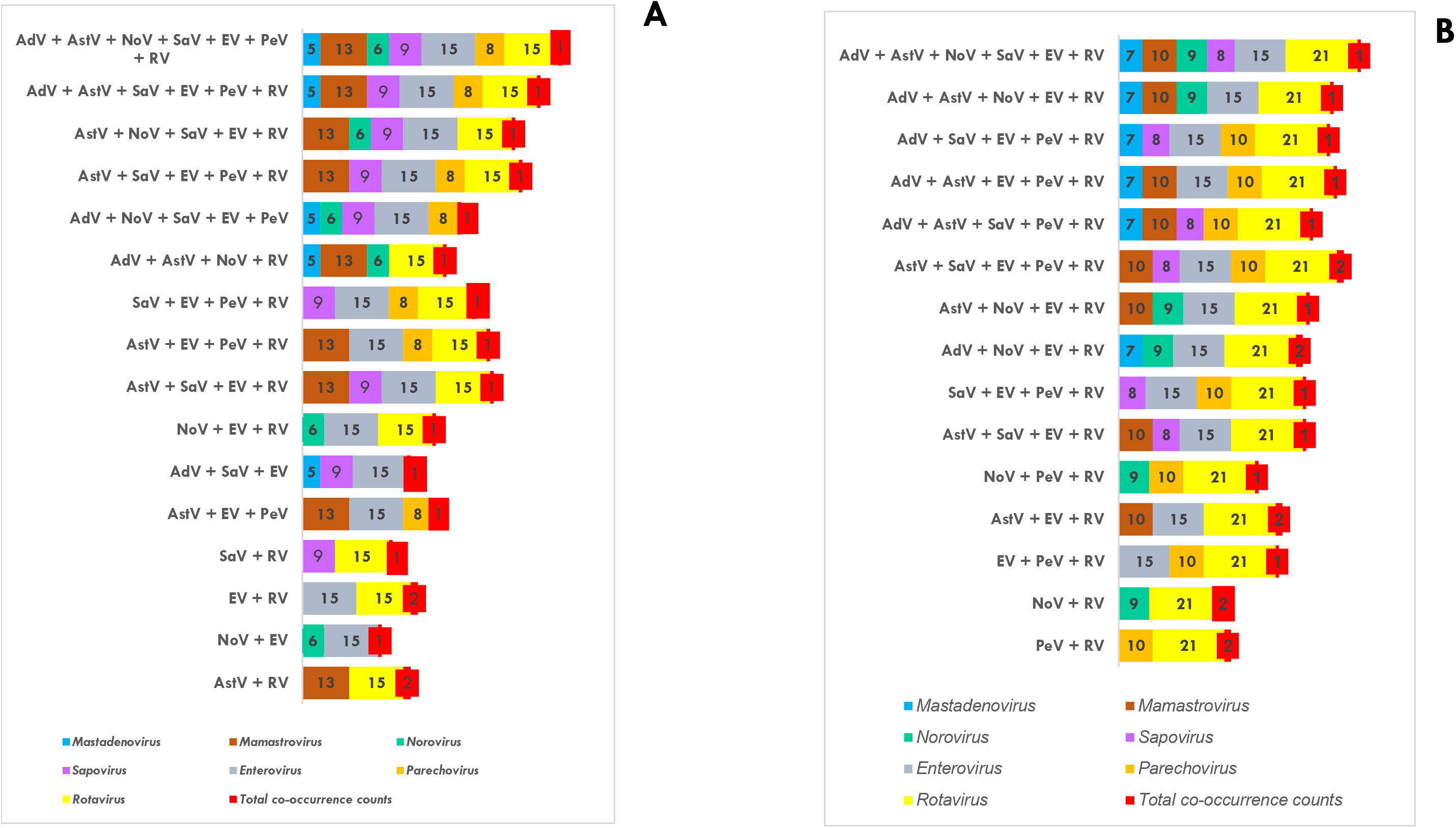
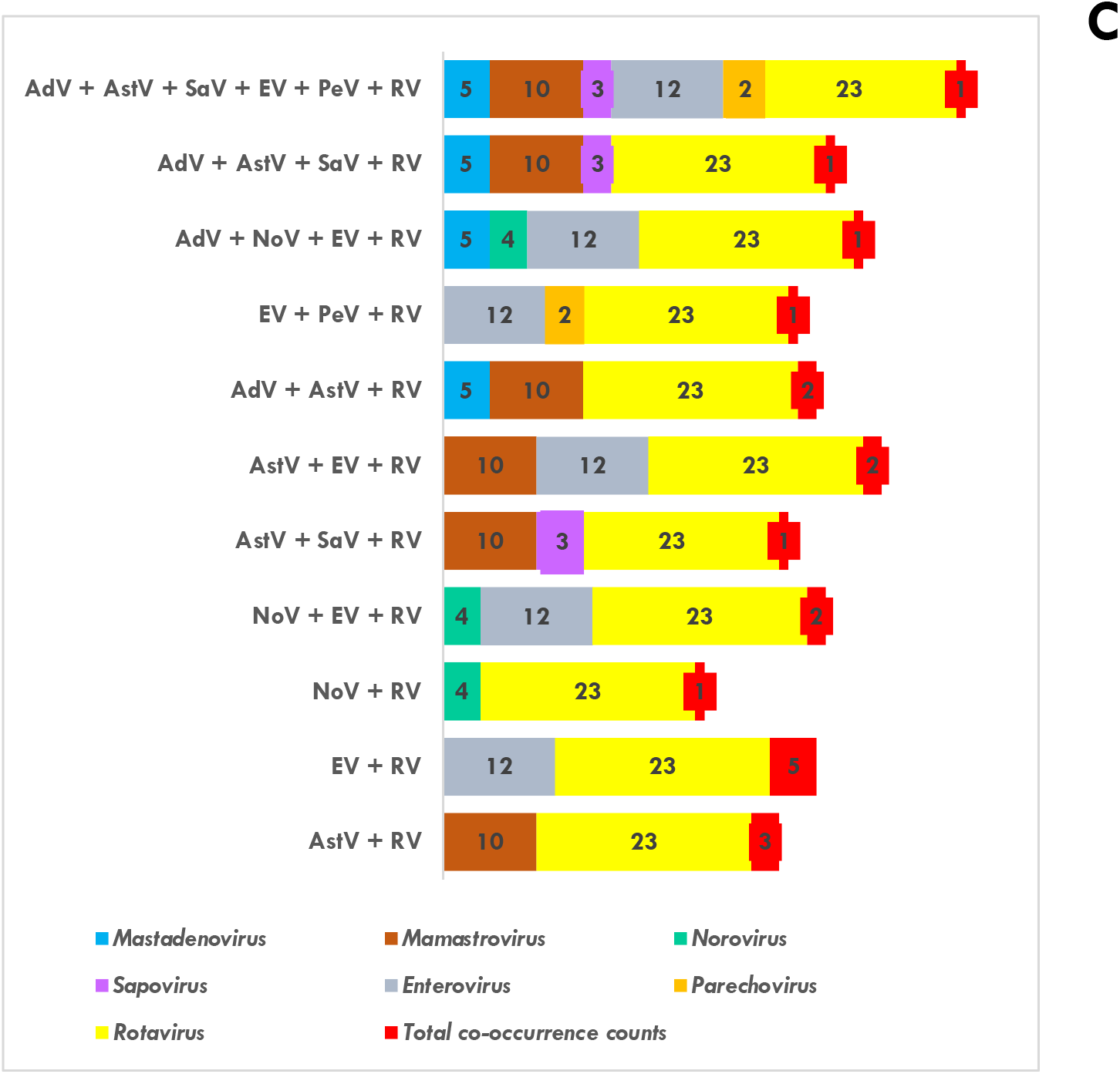
Distribution of co-detected enteric viral combinations detected in the study population across all three study cohorts: **(A)** Malnourished children with gastroenteritis (SAM), **(B),** Non-malnourished children with gastroenteritis (NAM), and **(C)** Children without gastroenteritis and malnutrition (NDM). Each horizontal staked bar represents one co-occurrence pattern of viral genera. Numbers within coloured segments indicate the detection frequency of each genus. Colours denote viral genera as follows: *Mastadenovirus* (HAdV) (blue), *Mamastrovirus* (HAstV) (brown), *Norovirus* (NoV) (green), *Sapovirus* (SaV) (purple), *Enterovirus* (EV) (grey), *Parechovirus* (PeV) (orange), *Rotavirus* (RV) (bright yellow), and in red, the overall co-occurrence counts for each pattern.

In the malnourished children with gastroenteritis (SAM), the most frequently detected genera among the co-occurring viruses were *Enterovirus* and *Rotavirus*, each detected in 15/21 samples, followed by AdV (13/21), SaV (9/21), PeV (8/21), Nov (6/21), and AstV (5/21). The SAM cohort demonstrated a broad range of co-occurrence patterns, including combinations involving two to seven viral genera. The most complex combination, comprising AdV, AstV, NoV, SaV, EV, PeV, and RV, was observed in one sample. Several other higher-order combinations involving five or more viral genera were also detected, although each occurred in relatively few samples. Among the pairwise combinations, EV + RV and AstV *+* RV were each observed in two samples, whereas most other combinations occurred in a single sample **(Fig. 7A).**

In the non-malnourished children with gastroenteritis (NAM), Rotavirus showed the highest detection frequency, occurring in 21/22 samples, followed by EV (15/22), AstV (10/22), PeV (10/22), NoV (9/22), AdV (7/22), and SaV (8/22). Multiple higher-order co-occurrence patterns were observed, with combinations involving up to six viral genera. Several combinations containing Rotavirus and multiple other enteric viruses were detected. Notably, AstV + SaV + EV + PeV + RV, AdV + NoV + EV + RV, and AstV + EV + RV each occurred in two samples, while most of the remaining combinations were detected in one sample. The predominance of Rotavirus within these combinations was consistent with its high detection frequency in the NAM cohort. Two dual viral co-detections were identified, NoV + RV and PeV + RV, with each combination detected in two samples. Triple viral co-occurrences were also observed **(Fig. 7B).**

Among children without gastroenteritis and malnutrition (NDM; control group), Rotavirus was detected in all 23/23 samples, followed by EV in 12/23 and AstV in 10/23 samples. AdV and NoV were detected in 5/23 and 4/23 samples, respectively, while SaV and PeV were detected less frequently, in 3/23 and 2/23 samples, respectively. Co-occurrence patterns in the NDM cohort were generally less complex than those observed in the two gastroenteritis cohorts, with combinations involving two to six viral genera. The most complex combination, comprising AdV, AstV, SaV, EV, PeV, and RV, occurred in one sample. Pairwise EV + RV co-occurrence was observed in five samples, representing the most frequent specific co-occurrence pattern in the control group and across the study groups, while AstV + RV occurred in three samples. Other combinations, including AdV + AstV + RV, AstV + EV + RV, and NoV + EV + RV, were observed at lower frequencies **(Fig. 7C).**

#### 3.5.8. Alpha diversity and Beta diversity of the infant gut virome across study groups

To explore and visualise virome diversity within the infant gut across the study cohorts, both alpha and beta diversity analyses were performed. Alpha diversity was assessed using the observed richness, Shannon, and Simpson diversity indices to evaluate within-sample viral diversity and richness. Beta diversity was evaluated using Principal Coordinates Analysis (PCoA), Bray-Curtis dissimilarity and Jaccard presence–absence distance matrices to assess differences in virome composition between study groups.

##### 3.5.8.1. Alpha diversity

The alpha diversity analysis based on the observed richness demonstrates differences in viral species richness among the three study cohorts (Fig. 8A). Non-malnourished children with gastroenteritis (NAM) exhibited the highest observed richness, followed by malnourished children with gastroenteritis (SAM). Children without gastroenteritis and malnutrition (NDM) displayed relatively lower and more homogeneous richness profiles, indicating a less complex virome composition. Further analysis of observed richness using the nonparametric Kruskal-Wallis test demonstrated a statistically significant difference (*p* = 0.0318) across the study groups. This indicates that the number of observed viral taxa differed significantly among the cohorts, suggesting variation in viral species richness within the infant gut virome. The wider distribution and higher median richness observed among children with gastroenteritis. When a pairwise comparison was made, significant results were obtained between NDM and NAM (*p* = 0.014); on the contrary, that between NDM and SAM and NAM and SAM showed no significance (*p* = 0.058 and *p* = 0.526), respectively, where *p* ˂ 0.05 is statistically significant **(Fig. 8A).**

**Fig. 8.**
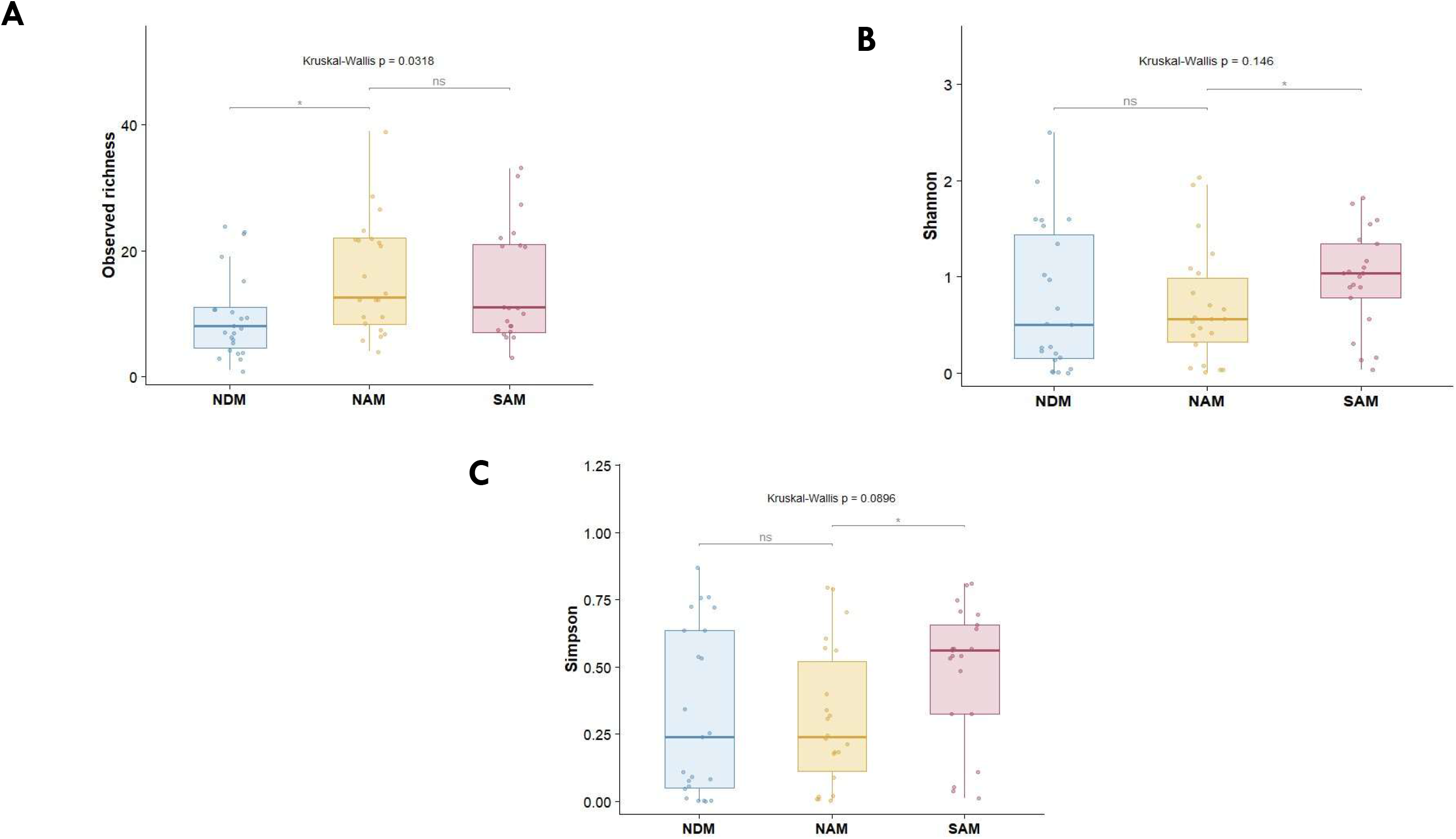
The alpha diversity metric illustrating the gut virome composition across the three study groups, each point represents a sample. NDM: Children without gastroenteritis and malnutrition; NAM: Non-malnourished children with gastroenteritis; SAM: Malnourished children with gastroenteritis. Each point represents an individual sample. **(A) Observed richness index (B) Shannon-Wiener index (C) Simpson index**. Statistically significant comparison difference was observed following Benjamini-Hochberg correction (BH) for observed viral richness between NDM and NAM (adjusted q = 0.042).

The Shannon index incorporates both richness and evenness, but measures the richness more within samples. Kruskal-Wallis’s statistics of the Shannon diversity analysis demonstrated no statistically significant difference, *p* = 0.1459, in overall virome diversity among the three study cohorts, namely NDM, NAM, and SAM **(Fig. 8B).** Despite the absence of overall statistical significance, distinct patterns were observed across the cohorts. The SAM group exhibited a higher median Shannon diversity than the NDM and NAM cohorts, suggesting a more even distribution of viral species within the gut virome of SAM children. In contrast, the NDM and NAM groups exhibited lower median diversity, suggesting that fewer viral taxa may have dominated the virome composition in these cohorts. The NDM cohort demonstrated substantial inter-individual variability, as reflected by the broad dispersion of Shannon index values and the presence of several high-diversity samples. This NAM cohort exhibited moderate variability, although most samples clustered within lower Shannon diversity ranges. Pairwise comparisons revealed no statistically significant difference between NDM and NAM cohorts (*p* = 0.796) and between NDM and SAM cohorts (*p* = 0.16), while statistical significance was observed between NAM and SAM groups (*p* = 0.047) **(Fig. 8B).**

Simpson’s index places greater emphasis on evenness of the viral species than richness alone. Analysis of the Simpson index revealed that the SAM cohort exhibited the highest median Simpson diversity, indicating a more even distribution of viral species and reduced dominance by a few taxa. Also, no single species dominated the virome in these children; instead, viral species were distributed more evenly across samples. It was also observed that samples from the SAM cohort were generally concentrated around higher Simpson values, suggesting a more uniformly distributed viral community among malnourished children with gastroenteritis. In contrast, both the NDM and NAM cohorts displayed lower median Simpson index values, suggesting that their viromes were characterised by greater dominance of a few highly abundant viral species. This observation is consistent with the predominance of certain enteric viruses such as Rotavirus, within these groups. The NDM cohort showed substantial variability in Simpson diversity, with values ranging from very low to relatively high, indicating considerable inter-individual heterogeneity in viral community structure among healthy children. Similarly, the NAM cohort exhibited broad variation, although most samples clustered at lower diversity values. The Simpson diversity index analysis revealed no statistically significant (Kruskal-Wallis, *p* = 0.0896), overall difference in virome diversity among the three cohorts **(Fig. 8C).** Pairwise Wilcoxon tests with Benjamini-Hochberg correction used to compare alpha-diversity metrics between the study cohorts revealed no statistically significant pairwise differences in Shannon diversity and Simpson diversity after correction for multiple testing (adjusted q-values ˃ 0.05). For observed viral richness, the comparison between NDM and NAM was statistically significant (adjusted q = 0.042).

##### 3.5.8.2. Beta diversity

The results of the beta diversity metrics across the three study groups were determined using principal coordinates analysis (PCoA) based on the Bray-Curtis and Jaccard indices. PCoA based on Bray-Curtis dissimilarity was performed to evaluate differences in virome composition among children across all three cohorts (SAM, NAM and NDM) **(Fig. 9A.).** The principal coordinates, PCoA 1 and PCoA 2 explained 15.2% and 12.0% of the total variation, respectively, accounting for 27.2% of the overall variation in virome composition. The ordination plot revealed substantial overlap among the three study groups, indicating that the overall virome composition was broadly similar across cohorts. The NDM samples were widely dispersed along PCoA 1, suggesting considerable inter-individual variability in viral community composition among healthy children. Similarly, NAM samples exhibited a broad distribution across both axes, with extensive overlap with the NDM group. In contrast, SAM samples appeared to form a relatively tight cluster, particularly along the PCoA 2, suggesting lower within-group variability and a more homogenous virome composition among malnourished children with gastroenteritis. Nevertheless, considerable overlap between SAM and the other cohorts was observed, indicating that the viral communities were not completely distinct. The PERMANOVA analysis on Bray-Curtis’s dissimilarity index revealed a statistically significant difference in virome composition among the three study cohorts (p=0.0038; R^2^=0.0563; df=2), despite the overlap. Pairwise comparison of PERMANOVA analysis on Bray-Curtis’s distance matrix revealed no significant statistical difference between NDM and NAM cohorts (R^2^=0.0285; *p*=0.202) and between NAM and SAM cohorts (R^2^=0.0373; *p*=0.0645), while statistical significance was observed between NDM and SAM groups (R^2^=0.0622; *p*=0.0015) (**Fig. 9A).**

**Fig. 9.**
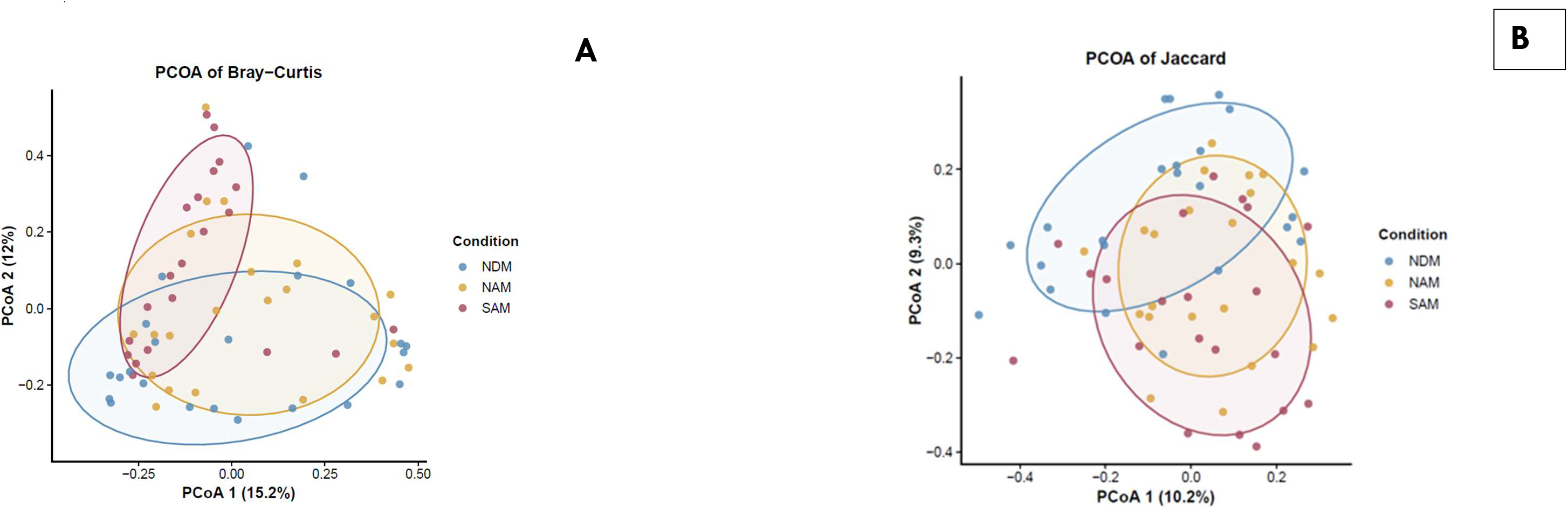
**A.** Principal coordinates analysis (PCoA) based on Bray–Curtis’s dissimilarity illustrating differences in gut virome composition among the three study cohorts. Differences in viral abundance-based community composition were statistically assessed using PERMANOVA (*p* = 0.0038). Each point represents an individual sample. NDM, children without gastroenteritis or malnutrition; NAM, non-malnourished children with gastroenteritis; SAM, malnourished children with gastroenteritis and **(B)** Principal coordinates analysis (PCoA) based on Jaccard dissimilarity, illustrating differences in gut virome composition based on viral presence/absence across the three study cohorts. Differences in virome composition among cohorts were assessed using PERMANOVA and were statistically significant (p = 0.0001). Each point represents an individual sample. NDM: children without gastroenteritis and malnutrition; NAM: non-malnourished children with gastroenteritis; SAM: malnourished children with gastroenteritis.

The Jaccard index is based solely on the presence or absence of taxa, making it less sensitive to quantitative shifts in viral populations than Bray-Curtis dissimilarity, an abundance-based metric. PCoA based on the Jaccard dissimilarity index was used to investigate differences in gut virome composition among the three study groups. The two principal coordinates explained 10.2% and 9.3% of the total variation, respectively, accounting for 19.5% of the overall variability in viral community composition **(Fig. 9B).** The ordination plot revealed substantial overlap among the three study groups, indicating that most of the viral taxa were shared across nutritional categories. Although NDM samples clustered toward higher values of PCoA 2, while SAM samples appeared more dispersed and shifted toward lower PCoA 2 values, no distinct separation between the groups was observed. NAM samples occupied an intermediate position and overlapped extensively with both NDM and SAM samples.

The PERMANOVA analysis on Jaccard dissimilarity index showed a statistically significant difference in virome composition among the three study cohorts (p=0.0001; R^2^=0.063149; df=2), despite the overlap. Pairwise comparison of PERMANOVA analysis on Jaccard diversity metrics revealed a significant statistical difference between NDM and NAM groups (R^2^=0.044588; *p*=0.00735) and between NDM and SAM cohorts (R^2^=0.064882; *p*=0.0003), while no statistical significance was observed between NAM and SAM groups (R^2^=0.034116; *p*=0.0594) **(Fig. 9B).**

### 3.6. Bacteriophage Composition of the Infant Gut Virome

In this study, bacteriophages were detected at a relatively low abundance (329,187 reads) compared with eukaryotic viruses, which accounted for 6.1% of the total viral reads. Of these bacteriophage-associated reads, 194,534 (59.1%) were detected in children without gastroenteritis and malnutrition, 87,840 (26.7%) in malnourished children with gastroenteritis, and 46,813 (14.2%) in non-malnourished children with gastroenteritis. Taxonomic classification of bacteriophage reads was limited: only 37,359 reads (11.35%) were assigned to 27 viral families, while the majority, 291,828 reads (88.65%), remained unclassified at the family level. Among the classified families, *Intestiviridae* was the most abundant, with 12,580 reads accounting for 3.82% of the total bacteriophage reads. This was followed by *Drexlerviridae* and *Microviridae* which represented of 2.97% and 1.05% of bacteriophage reads, respectively. The remaining families were detected at substantially very lower abundances (**Fig. 10A**). Apart from *Microviridae* and *Fiersviridae*, the classified bacteriophages were predominantly crAss-like phages, which belong to the order of *Crassvirales* within the class *Caudoviricetes*. CrAss-like phages are among the most abundant and widely distributed bacteriophages in the human gut currently known as *Crassvirales* within the class *Caudoviricetes,* which are the most abundant phages in the human gut.

**Fig. 10A.**
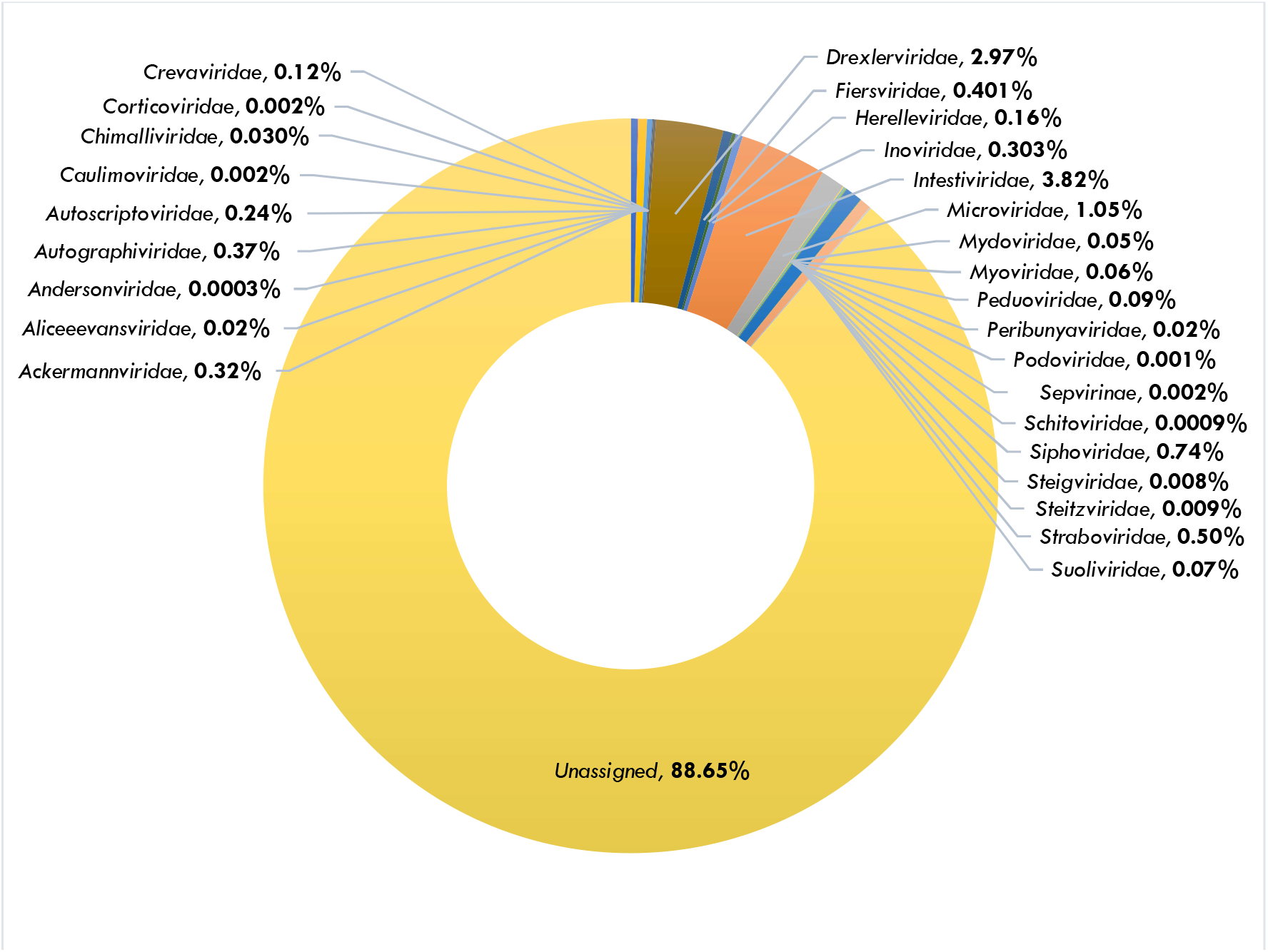
Comparative taxonomic distribution and read abundance of bacteriophage families across malnourished children with gastroenteritis, non-malnourished children with gastroenteritis, and children without gastroenteritis and malnutrition.

**Fig. 10B.**
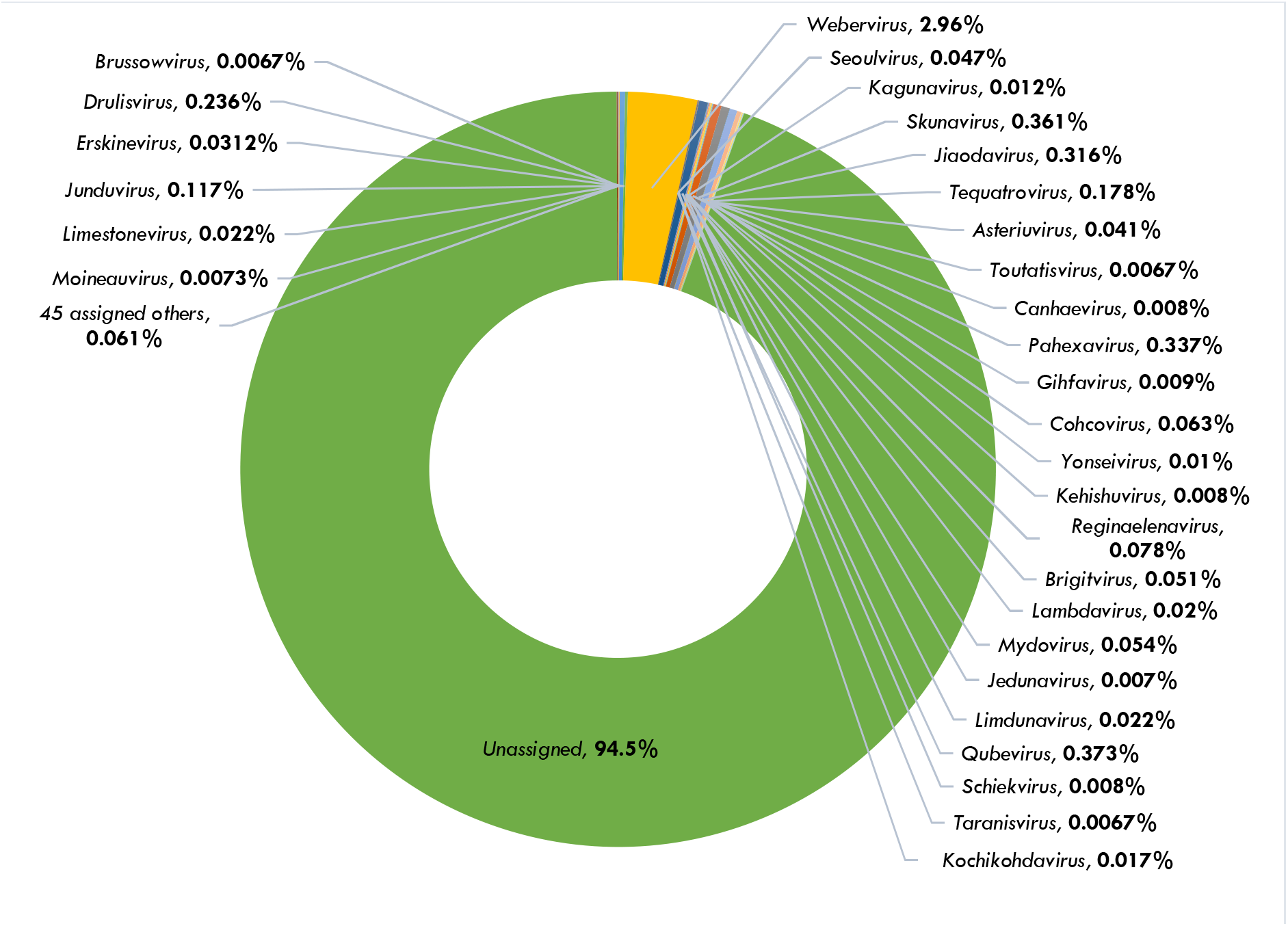
Relative proportions and taxonomic distribution of bacteriophage genera identified from sequencing reads in infant faecal samples across the three-study cohort; malnourished children with gastroenteritis, non-malnourished children with gastroenteritis, and children without gastroenteritis and malnutrition.

At the genus level, taxonomic assignment was similarly limited, with 311,185 (94.5%) of the reads remaining unclassified, and only 18,033 reads (5.5%) could be assigned to seventy-five genera. The predominant genus was Webervirus, belonging to the family Drexlerviridae, with 9,754 reads, accounting for 2.96% of the total phageome reads. Other detected genera included *Qubevirus* (0.373%), *Skunavirus* (0.361%), *Pahexavirus* (0.337%), *Jiaodavirus* (0.316%), *Drulisvirus* (0.236%). The remaining genera were represented by comparatively very low reads across all study cohorts. Overall, crAssphages dominated the infant gut phageome in all three study cohorts (**Fig. 10B**), suggesting that these viruses constitute a core component of the infant gut virome regardless of gastroenteritis or nutritional status.

## 4.0. Discussion

Enteric viruses comprise a diverse group of viral pathogens that are major aetiological agents of gastroenteritis worldwide and can be detected in both symptomatic and asymptomatic children under five years of age (Liang et al., 2022; Okitsu et al., 2020). Advances in viral metagenomics, particularly mNGS, have greatly improved the comprehensive profiling of the infant gut virome and the identification of viral pathogens associated with infectious diseases, especially in cases where the causative agent is unknown (do Socorro Fôro Ramos et al., 2021; Fernandez-Cassi et al., 2020; Mohammad et al., 2020).

Of the 69 samples sequenced, 66 yielded a large proportion of viral sequence data for downstream analyses. The predominance of eukaryotic viral reads is notable, given that bacteriophages are widely recognised as abundant components of the human gut virome (Fulci et al., 2021). This finding is consistent with that reported in previous metagenomic studies conducted in South Africa (Milton Tshidiso Mogotsi et al., 2024), and Brazil (do Socorro Fôro Ramos et al., 2021), but contrasts with reports of bacteriophage predominance in the gut virome (Koonin and Yutin, 2020; Yinda et al., 2019). The high proportion of eukaryotic viral reads may partly reflect the substantial contribution of enteric viruses, particularly Rotavirus A and Sapovirus, detected in this study. Differences between studies may also be attributed variations in study populations, age groups, geographical settings, sample processing protocols, viral enrichment strategies, sequencing approaches and bioinformatic pipelines. In the present study, viral enrichment using sequential filtration and nuclease treatment substantially reduced host and bacterial nucleic acids, thereby increasing the proportion of viral reads available for metagenomic analysis ( Hall et al., 2014; Cotten et al., 2014).

The observed highest abundance of eukaryotic viral reads (37.2%) for non-malnourished children with gastroenteritis highlights the substantial representation of eukaryotic viruses in children with gastroenteritis in the absence of malnutrition. It may reflect differences in the distribution and relative abundance of viral taxa among the study cohorts, particularly the predominant enteric viruses which are important causes of childhood gastroenteritis (Omatola et al., 2024). The predominant enteric viruses detected included members of the genera *Mastadenovirus (MAdV)*, *Mamastrovirus (MAstV)*, *Norovirus (NoV)*, *Sapovirus (SaV)*, *Enterovirus (EV)*, *Cardiovirus (CaV)*, *Cosavirus (CoSV)*, *Parechovirus (PeV)*, and *Rotavirus (RV)*, all of which have previously been associated with childhood gastroenteritis in sub-Saharan Africa (Oppong et al., 2020; Yinda et al., 2019), South Africa (Mafokwane et al., 2023), and globally (Yang et al., 2022; Xavier et al., 2020). *RV* predominated, and as the leading aetiological agent of severe viral gastroenteritis in children under five years of age worldwide. Rotavirus may have contributed to the virome observed in this cohort but read abundance alone does not establish causation. Nevertheless, the co-detection of other established enteric viruses suggests that multiple viral pathogens may also have contributed to the clinical presentation.

Healthy children without gastroenteritis and malnutrition accounted for 32.3% of total eukaryotic reads, representing the second most abundant cohort. Similar to the gastroenteritis cohorts, the predominant enteric viruses belonged to the genera *AdV*, *AstV*, *NoV*, *SaV*, *EV*, *PeV*, and *RV*. *RV* was overwhelmingly dominant, representing 96.1% of the reads, with more than one million reads detected in a single asymptomatic child. This high abundance most likely reflects vaccine-derived RV shedding following administration of the oral Rotarix vaccine, although prolonged viral shedding after vaccination, recent asymptomatic infection, or exposure to vaccine strains transmitted from recently vaccinated children cannot be excluded. Likewise, the detection of *EV* may reflect shedding following oral poliovirus vaccination. Despite the high abundance of enteric viruses, none of the children in this cohort exhibited gastrointestinal symptoms or evidence of malnutrition. Similar observations have been reported in Brazil, South Africa, and Bangladesh, where *RV*, *NoV*, *SaV*, *AstV*, *EV*, and *PeV* were frequently detected in asymptomatic children (Mogotsi et al., 2024; Rivera-Gutiérrez et al., 2023; Okitsu et al., 2020). Collectively, these findings suggest that asymptomatic carriage of enteric viruses during early childhood is common, although it remains to be determined whether this reflects prolonged viral shedding, persistent infection, repeated exposure, or vaccine-derived virus.

Lastly, the cohort of malnourished children with gastroenteritis accounting for 30.4% of all classified eukaryotic viral reads. This finding may reflect lower viral shedding at the time of sampling and the relatively smaller sample size. A broad diversity of enteric viruses was identified, including members of the genera M*AdV*, M*AstV*, *NV*, *SaV*, *CoSV*, *EV*, *PeV,* and *RV*. In addition, *Kobuvirus* and *Salivirus* were detected; however, their roles in gastroenteritis and malnutrition remain poorly understood and require further investigation (do Socorro Fôro Ramos et al., 2021; Yu et al., 2015). Unlike the other two cohorts, *SaV* was the predominant enteric virus, followed by *EV*. In contrast, *RV* had a lower abundance than *SaV* and *EV*. Notably, *RV* read abundance increased progressively from malnourished children with gastroenteritis to non-malnourished children with gastroenteritis, and then to healthy controls. This lower reads abundance in malnourished children may be explained by the intestinal pathology associated with malnutrition, including villous atrophy and the reduced availability of mature enterocytes required for efficient rotavirus replication (Burnett et al., 2021). Malnutrition is also associated with impaired immune function, altered gut microbial and viral communities, prolonged enteric infections, and an increased frequency of co-infections, all of which may influence viral composition and disease severity (Reyes et al., 2015).

In addition to established human enteric viruses, a wide range of non-enteric human viruses and viruses associated with animals, plants, fungi, protists, and birds were detected across the three study cohorts. The presence of these viruses most likely reflects dietary intake, environmental exposure, transient passage through the gastrointestinal tract, or asymptomatic carriage rather than active infection of the intestinal tract (Pargin et al., 2023; Tan et al., 2020; McCann et al., 2018). Collectively, these findings demonstrate that the infant gut virome is a highly diverse and complex ecosystem, irrespective of the presence or absence of gastroenteritis and malnutrition.

Recently, metagenomic studies have reported *Caliciviridae* and *Picornaviridae* as the predominant enteric viral families based on both read abundance and detection frequency (Rivera-Gutiérrez et al., 2023; Yang et al., 2022). In contrast, the current study identified *Sedoreoviridae* and *Caliciviridae* as the predominant enteric viral families, together accounting for 83.09% of the total eukaryotic viral reads.

*Rotavirus A* (RVA), the only species within the genus Rotavirus of the family *Sedoreoviridae,* was the most abundant enteric virus, representing 59.53% of all eukaryotic viral reads and was detected across all three study cohorts. RVA remains the leading cause of severe viral gastroenteritis in children under five years of age, particularly in sub-Saharan Africa and Southeast Asia (Omatola et al., 2024; Tian et al., 2024; Oppong et al., 2020; Rossouw et al., 2021). Infection is often asymptomatic or mild in young infants because of maternally acquired antibodies, whereas severe disease can result in dehydration, electrolyte imbalance, shock, and death (Greenberg and Estes, 2009). Although, the children in this study had received the rotavirus vaccine, RVA was detected in 89.4% of the study participants. This observation may reflect continued shedding of vaccine-derived virus, infection with circulating wild-type strains, or breakthrough infection with genotypes that differ from those represented in the vaccine (Mwape et al., 2023). However, because viral genotyping was beyond the scope of this study, these possibilities could not be distinguished. Continued genomic surveillance is therefore essential to monitor the circulation and evolution of rotavirus strains in South Africa. The continued predominance of RVA observed in our study is consistent with previous reports from Africa describing the emergence and circulation of diverse rotavirus genotypes, emphasizing the importance of ongoing molecular surveillance to evaluate vaccine performance and monitor genotype replacement (Page et al., 2018; Gómez et al., 2014; Seheri et al., 2012).

*Rotavirus C* (RVC) was detected at a low abundance (21 reads), predominantly in non-malnourished children with gastroenteritis and healthy controls. RVC was first identified during an outbreak in Ohio, USA, in 1980 (Saif et al., 1980) and has since been reported in humans as well as several animal species, including pigs, cattle, dogs, ferrets, and minks (Xie et al., 2018; Tiku et al., 2017; Marton et al., 2015; Mawatari et al., 2014). Pigs are considered the principal reservoir of RVC, and sporadic porcine-to-human transmission events have been reported in Brazil and India (Bhat et al., 2018; Gabbay et al., 2008). Although the source of RVC detected in this study could not be determined, its presence may reflect environmental exposure or possible zoonotic transmission, particularly in regions where pig farming is practiced, including parts of KwaZulu-Natal (Gcumisa, 2013). Previous studies have also documented RVC infections in children from Kenya (Mwenda et al., 2003), Japan (Kumazaki and Usuku, 2014), South Korea (Tiku et al., 2017), and India, where RVC was associated with outbreaks of acute gastroenteritis (Joshi et al., 2017).

In addition to human rotaviruses, animal-associated rotaviruses, including *Bat rotavirus*, *Equine rotavirus*, and *Feline rotavirus,* were detected at low read abundances. *Bat rotavirus* was identified in a single non-malnourished child with gastroenteritis, while *Equine rotavirus* was detected across all three study cohorts in seven samples. *Feline rotavirus* was detected exclusively in non-malnourished children with gastroenteritis. Similar rotaviruses have previously been reported in bats, horses, and cats (Nakagomi and Nakagomi, 2026; Eertink et al., 2024; Yinda et al., 2016). Although, these viruses are primarily associated with their respective animal hosts, their detection in infant faecal samples may reflect environmental contamination, dietary exposure, or potential interspecies transmission. However, further genomic characterization and epidemiological investigations are required to determine their origin and assess their zoonotic significance.

*Caliciviridae* family was the second most abundant family in this current study. *Sapovirus* (SaV) and *Norovirus* (NoV) are both genera within the *Caliciviridae* family, with SaV being the dominant contributor to the high abundance recorded, with over a million reads. SaV was the second most abundant enteric virus after RVA detected in this study. SaV was detected in all cohorts with the highest detection being in the cohort consisting of malnourished children with gastroenteritis, and over a million reads were detected in a single sample within this cohort, probably due to recent infection. The reads abundance of SaV was much lower in non-malnourished children with gastroenteritis and in the control group. The same pattern was also observed at the level of the detection frequencies. Overall, SaV was detected in 20/66 (30.30%). SaVs cause acute gastroenteritis (AGE) in humans and animals (Oka et al., 2015), and has been associated with both sporadic cases and outbreaks among children, older adults, and immunocompromised individuals (de Oliveira-Tozetto et al., 2021; Becker-Dreps et al., 2020; Varela et al., 2019). SaV can be asymptomatically shed in children, including those without AGE symptoms (Rivera-Gutiérrez et al., 2023; Pitkänen et al., 2022).

Following the widespread introduction of rotavirus vaccines, the relative contribution of other enteric viruses, including SaV, has become increasingly important (Carvalho et al., 2025). SaV prevalence in low- and middle-income countries is estimated at 6.5%, and it is recognized as an important viral cause of gastroenteritis, after RV and NoV, (Inns et al., 2019). Murray et al. detected SaV in faecal samples collected from children younger than six years old hospitalised with gastroenteritis over a five-year period from April 2009 to December 2013, in the Gauteng, KwaZulu-Natal, Mpumalanga and Western Cape provinces of South Africa. SaV single infections were present in 57% of the specimens from which SaVs were genotyped (Murray et al., 2016). This prevalence was higher than that observed in our study. Conversely, Mogotsi et al. identified SaV as the most abundant enteric virus among asymptomatic children in the Free State Province, South Africa (Mogotsi et al., 2024). SaV has also been reported among children with gastroenteritis in several African countries, including Burkina Faso (Matussek et al., 2015), Tunisia, (Sdiri-Loulizi et al., 2011), Tanzania (Liu et al., 2011) and in Malawi (Dove et al., 2005). Similarly, in Jiangsu China, SaV was detected in 19% of children with AGE with unknown aetiology (Li et al., 2020).

*Norovirus* (NoV), another member of the *Caliciviridae* family, has become the leading cause of epidemic and sporadic AGE worldwide following the introduction of rotavirus vaccination programmes (Yang et al., 2022). The highest abundance of NoV was observed in malnourished children with gastroenteritis, followed by non-malnourished children with gastroenteritis, and healthy controls. The higher NoV abundance observed in malnourished children with gastroenteritis may indicate increased viral burden or prolonged shedding in this vulnerable group. Given the capacity of NoV to undergo prolonged shedding in symptomatic and asymptomatic individuals, this finding may have implications for infection persistence and transmission (Tk et al., 2019; Ramani et al., 2016). However, the detection of NoV alone does not establish it as the cause of gastroenteritis, particularly in the presence of asymptomatic shedding or mixed infections. A study in Brazil detected NoV in 45.2% of children with diarrhoea (Gondim et al., 2018). A metagenomic study conducted by Rivera-Gutiérrez et al identified NoV as one of the predominant enteric viruses in healthy Mexican infants (Rivera-Gutiérrez et al., 2023). Chukwu et al., reported NoV predominantly among children younger than two years in South Africa (Chukwu et al., 2020). Although NoV infections are generally self-limiting, they can cause severe dehydration requiring hospitalisation in young children. Currently, no licensed vaccine is available, although several virus-like particle-based vaccine candidates are undergoing clinical development (Esposito and Principi, 2020).

In this study, five human adenovirus (HAdV) species (HAdV-A, HAdV-B, HAdV-C, HAdV-D, and HAdV-F) were detected. The predominance of HAdV-F observed in the present study reinforces the well-established role of enteric adenovirus species F (types 40 and 41) as important causes of paediatric gastroenteritis (Yodmeeklin et al., 2023). These findings are consistent with previous reports identifying HAdV-F as the predominant adenovirus associated with childhood diarrhoeal disease, although the reported prevalence varies considerably between geographical regions. For example, Li et al. (2020), reported a lower HAdV detection rate of 3.1% among children with acute diarrhoea in Hangzhou, China, whereas Tahmasebi et al. (2020), reported a substantially higher prevalence of 57.8% among children with acute gastroenteritis in Northern Brazil using next-generation sequencing. Similarly, Gelaw and Liebert (2022) detected HAdVs in 18.4% of children presenting with acute gastroenteritis in Northwest Ethiopia, a slightly lower prevalence than that observed in the present study. The differences in the prevalence data could be due to different laboratory methods applied, different enrichment procedures, and sequencing depths. Furthermore, the data obtained regarding HAdV-F are consistent with those of Potgieter et al. (2023), who reported HAdV-F (types 40/41) as the predominant adenovirus among South African children under five years of age with diarrhoea in Limpopo Province, further supporting its important role in paediatric gastroenteritis in South Africa. Although, HAdV-F is the primary enteric adenovirus associated with gastroenteritis, adenoviruses are also known to infect the respiratory tract and may subsequently be shed in faeces, even in the absence of gastrointestinal disease (Xu et al., 2025). This may explain the detection of non-enteric HAdV species across all study cohorts and highlights the importance of interpreting adenovirus detection in faecal samples within the broader clinical context.

In this present study, *Astroviridae* was detected across all study groups and was the fourth most detected enteric eukaryotic viral family, with a detection rate of 50.0%, with the highest reads being recovered from non-malnourished children with gastroenteritis. MAstVs 1, 2, 3, and 8 were detected, with MAstV 1 predominating, confirming its role as the predominant circulating human astrovirus. Similarly, the Malnutrition and Enteric Disease Study (MAL-ED) reported astrovirus infection in 35% of children and demonstrated that malnutrition, particularly stunting, is a risk factor for astrovirus-associated diarrhoea (Olortegui et al., 2018). However, the comparatively low abundance of astroviruses in malnourished children with gastroenteritis in our study suggests that they were unlikely to be the predominant viral agents responsible for disease in this cohort, although their contribution cannot be excluded. Further studies are therefore needed to clarify the relationship between astrovirus infection, gastroenteritis, and malnutrition.

Novel human astroviruses were also identified, including MAstV 8, also known as *Human-Mink-Ovine Astrovirus A* (HMOAstV-A), which has previously been reported in humans and sheep from Egypt, Japan, United States of America, Kenya, and China (Vu et al., 2016). Animal-associated astroviruses, including MAstV-2, also known as Feline astrovirus (FeAstV), MAstV-3, for *Porcine astrovirus* (PoAstV), *Bovine astrovirus* (BoAstV), and *California sea lion astrovirus 11,* were also detected at relatively low abundance. Their presence supports previous evidence that interspecies transmission of astroviruses may occur, highlighting that the detected sequences may warrant investigation of host range, without establishing zoonotic transmission (Bosch et al., 2014). Furthermore, the detection of astroviruses in both symptomatic and asymptomatic children is consistent with findings in a study from South Africa (Khumela et al., 2021)and highlights the considerable genetic diversity of astroviruses circulating in paediatric populations.

*Picornaviridae* was the second most frequently detected enteric viral family in this study, with a prevalence of 84.9%. A wide diversity of both enteric and non-enteric picornaviruses was identified, with species of the genus *Enterovirus* (EV) predominating, accounting for above 60% of the total reads assigned to picornaviruses. The prevalence of EV infections in children with gastroenteritis has been reported to range from 1.2% to 42% worldwide (Shen et al., 2019), while asymptomatic infections (Rao, 2021) and co-infections with other enteric viruses have also been documented (Tatte and Gopalkrishna, 2019). In the present study, Enterovirus A, B, C, D, F, G, and J were detected. Among the human enteroviruses, *Enterovirus B* (EVB) was the most abundant, and was detected at 18.18%, exclusively in the gastroenteritis cohorts, with the highest abundance observed in non-malnourished children with gastroenteritis. Although a higher prevalence of 32% has been reported in China (Rojjanadumrongkul et al., 2020), our findings demonstrate its circulation among South African children with gastroenteritis. Coxsackievirus and Echovirus, both viral species associated with gastroenteritis, were detected in non-malnourished children with gastroenteritis. *Enterovirus C* (EVC) was the most frequently detected enterovirus, occurring in 25/66 (37.88%), predominantly in the control group. This finding may reflect shedding of the oral poliovirus vaccine (OPV), which forms part of South Africa’s childhood immunisation programme, although vaccination records were not available in this study. The widespread use of OPV has contributed substantially to global polio eradication, despite continued challenges posed by circulating vaccine-derived polioviruses (Jartti et al., 2024).

Human rhinoviruses (HRV) A, B, and C, which are common causes of respiratory tract infections (Yang et al., 2022; Martin et al., 2018), were also detected at relatively low abundance, with HRV-A being the predominant species. Since respiratory specimens were not collected, it was not possible to determine whether these viruses originated from swallowed respiratory secretions or true gastrointestinal infection. Their detection is nevertheless consistent with a previous report demonstrating that respiratory viruses can survive gastrointestinal passage and be shed in faeces (Xu et al., 2025). Therefore, their presence in stool should not be interpreted as evidence that they are the causative agents of gastroenteritis.

*Parechovirus* (PeV), another genus within the *Picornaviridae* family associated with gastroenteritis in children under five years of age, has a reported global prevalence ranging from 1.8% to 29.4% (Yang et al., 2022). In the present study, PeV was detected across all three study cohorts at 30.3%, a detection frequency slightly higher than the reported global prevalence. Their detection does not establish a causal role in gastroenteritis, particularly because these viruses were also detected in asymptomatic children. Three HPeV species, HPeV1, HPeVA, and HPeV sp., were identified. Although HPeV1 has been reported as the predominant species, HPeVA was the most frequently detected and abundant species in this study. Furthermore, the detection of HPeV in asymptomatic children is consistent with previous reports indicating that *Parechovirus* infections are often asymptomatic (Mogotsi et al., 2024; Zhang et al., 2022).

*Cosavirus* (CoSV) is a genus within the *Picornaviridae* family that consists of several human enteric viruses that have been associated with gastroenteritis in humans. They were first identified in stool samples from South Asian children with non-polio acute flaccid paralysis (Kapoor et al., 2008), and have subsequently been reported at relatively low prevalence in Asia, Australia, Africa, Europe, North and South America (Khamrin and Maneekarn 2014; Okitsu et al. 2014; Rovida et al. 2013; Kapoor et al. 2008; Dai et al., 2010). In the present study, CoSV was detected among malnourished and non-malnourished children with gastroenteritis at a prevalence of 21.12%, while no detections were observed in the control group. Two species, CoSV A and CoSV D, were identified. A considerably higher prevalence (41.4%) was previously reported among children in rural Bangladesh, where CoSV was also detected in asymptomatic children (Okitsu et al., 2020), unlike our findings. Similarly, a longitudinal study in Ghana identified a novel CoSV A strain in asymptomatic children and suggested that it may reduce the immunogenicity of rotavirus and oral poliovirus vaccines (Kim et al., 2022). Despite increasing reports of CoSV detection, its pathogenic role in diarrhoeal disease remains uncertain and warrants further investigation (Okitsu et al., 2020).

*Cardioviruses* (CaVs), members of the *Picornaviridae* family, were first isolated in rodents (Pritchard et al., 1992). In 2007, a novel human cardiovirus, identified as scaffold (SAFV) was detected for the first time in a faecal sample of a child with fever, although its pathogenic role remains incompletely understood (Jones et al., 2007; Ugai et al., 2019). Three CaV species (A, B, and C) have traditionally been recognised, with more than 11 genotypes of CaV B associated with human infections, particularly SAFV-2 and SAFV-3 (Xavier et al., 2020). In this present study, CaV was detected exclusively in non-malnourished children with gastroenteritis. *CaVs A, B, and D* were the species detected in this present study, and *CaV B* was predominant, supporting previous reports of its association with paediatric gastroenteritis. Although SAFVs have been detected in stool samples of children with gastroenteritis as well as in environmental and sewage samples, they are not routinely included in clinical diagnostic testing (Vandesande et al., 2020; Bonanno Ferraro et al. 2019; Fernandez-Cassi et al., 2018). Consequently, their contribution to the burden of viral gastroenteritis remains uncertain and warrants further investigation.

Members of the *Parvoviridae* family are small, non-enveloped viruses with single-stranded DNA genomes that infect both vertebrate and invertebrate hosts and are classified into the subfamilies *Parvovirinae* and *Densovirinae*, respectively (Cotmore et al., 2019). In the present study, *Human bocavirus* (HBoV), a member of the genus *Bocavirus* within the *Parvovirinae* subfamily, was detected in both the malnourished and non-malnourished cohorts of children with gastroenteritis. HBoV was identified in two samples, yielding a combined total of 5083 reads. Although HBoV has been associated with respiratory and gastrointestinal infections, its pathogenic role in gastroenteritis remains uncertain (Khamrin et al., 2012). Four HBoV genotypes (HBoV1–4) have been identified in respiratory and faecal samples from children (Arthur et al., 2009; Kapoor et al., 2009). Therefore, the detection of HBoV in this study does not allow a definitive conclusion regarding its association with gastroenteritis or malnutrition. Members of the genus *Dependoparvovirus*, including *Adeno-associated virus (AAV)* and *Dependoparvovirus primate 1*, were the first parvoviruses identified in humans and are known to cause any symptoms (Liang et al., 2022). AAVs are replication-defective viruses that depend on coinfection with helper viruses, such as adenoviruses or herpesviruses, to support productive replication (Hildebrandt et al., 2020; Phan et al., 2012). Although AAVs can infect humans, they have not been definitively established as causes of human disease (Hildebrandt et al., 2020). In addition, *Bufavirus-2*, a member of the genus *Protoparvovirus*, was detected in the cohort consisting of non-malnourished children with gastroenteritis, and has previously been reported in the faeces of a child with acute diarrhoea in Burkina Faso (Phan et al., 2012). Other detected parvoviruses included members of the genera *Aveparvovirus*, *Copiparvovirus*, and *Chaphamaparvovirus*, as well as several unclassified members of the *Parvoviridae* family.

In addition to *Adenoviruses,* several DNA viruses belonging to the *Anelloviridae* family were detected across all three study cohorts. These included *Torque teno virus* (TTV) and *Alphatorquevirus homin 13* (genus *Alphatorquevirus*), *Torque teno mini virus* (TTMV; genus *Betatorquevirus*), *Gammatorque teno midi virus* (genus *Gammatorquevirus*), and *Gyrovirus chicken anemia* and *Gyrovirus galga 1* (genus *Gyrovirus*). Members of the *Anelloviridae* family were most abundant in the cohort of malnourished children with gastroenteritis. Our findings are consistent with those of Reyes et al., who reported an increased presence of A*nelloviruses* in malnourished discordant twin pairs, further supporting the association between A*nellovirus* abundance and malnutrition (Reyes et al., 2015). However, these viruses have not been directly implicated in the aetiology of gastroenteritis and are frequently detected in healthy, asymptomatic children (Liang et al., 2020; Maqsood et al., 2019; Focosi et al., 2016). Lim et al. reported that *Anelloviruses* become the predominant eukaryotic viruses of the infant gut virome by two years of age, likely reflecting continuous environmental exposure and early-life colonization (Lim et al., 2015). Consistent with their ubiquitous distribution, *Anelloviruses* have been identified at multiple body sites and have not been conclusively associated with any specific clinical disease (do Socorro Fôro Ramos et al., 2021). Although increased *Anellovirus* abundance has been reported in immunocompromised individuals, including patients living with HIV (Liang and Bushman, 2021), these viruses were detected in both children living with and without HIV in the present study, suggesting that their presence is unlikely to be directly associated with HIV status or gastroenteritis.

Coronaviruses are enveloped, positive-sense RNA viruses belonging to the *Coronaviridae* family and are primarily associated with respiratory disease in humans and a wide range of animal hosts, although different organ systems are also affected with varying severity (Li et al., 2019). Bats, particularly Chinese horseshoe bats, are considered the natural reservoir of several severe acute respiratory syndrome-related coronaviruses (SARS-CoV), with zoonotic transmission to humans occurring either directly or through intermediate animal hosts (Lau et al., 2005; Li et al., 2005; Guan et al., 2003). In the present study, *Betacoronavirus 1* was detected in a single sample from a malnourished child with gastroenteritis, yielding 3,205 reads. In contrast, *Betacoronavirus severe acute respiratory syndrome-related coronavirus* was identified in three non-malnourished children with gastroenteritis, totalling 51 reads. Both viral species belong to the genus *Betacoronavirus*. Although gastrointestinal symptoms, including diarrhoea, have been reported in coronavirus infections, they occur in only a proportion of infected individuals and are generally secondary to respiratory disease. The low abundance and limited detection of these respiratory-associated coronaviruses suggest that they were minor components of the faecal virome in the study population; therefore, they do not establish them as the aetiological agents of gastroenteritis. Their presence may instead reflect gastrointestinal shedding following respiratory infection, environmental exposure, dietary contamination, or transient passage through the gastrointestinal tract (O’Shea et al., 2019).

Human metapneumovirus (HMPV; *Metapneumovirus hominis*), a member of the genus *Metapneumovirus* within the family *Pneumoviridae*, was detected at a very low read abundance in the stool sample of a non-malnourished child with gastroenteritis. HMPV is a well-established cause of upper and lower respiratory tract infections, particularly in young children, older adults, and immunocompromised individuals, with the greatest disease burden occurring in children under five years of age (Feuillet et al., 2012). HMPV is known to circulate in South Africa and has been identified as an important respiratory pathogen among young children, particularly those living with HIV (Groome et al., 2015). Although HMPV RNA has occasionally been detected in faecal samples, there is currently no convincing evidence to support a direct role for HMPV in the pathogenesis of AGE.

A diverse range of plant-infecting viruses has been detected in the human gut virome, including that of infants (Yan et al., 2021; Zhang et al., 2006). In the present study, sequence reads corresponding to plant viruses that are not directly associated with gastroenteritis or malnutrition were identified across all three study cohorts, with the family *Virgaviridae* predominating. The genus *Tobamovirus* was represented by *Pepper mild mottle virus* (PMMoV), *Rehmannia mosaic virus* (RMV)*, Tobacco mild green mosaic virus* (TMGMV), *Tobacco mosaic virus* (TMV), *Tomato mosaic virus* (ToMV), *Tomato mottle mosaic virus* (ToMMV), and *Cucumber green mottle mosaic virus* (CGMMV), with TMGMV being the most abundant species. Similar findings were reported by Yan and colleagues who detected *Virgaviridae* members in the gut viromes of healthy Chinese individuals and visiting healthy Pakistanis, with viral composition associated with dietary habits, alcohol consumption, and smoking (Yan et al., 2021). Our findings are consistent with those of Mogotsi et al. (2024), who reported *Virgaviridae* as the predominant plant viral family in faecal samples from healthy South Africa children. Additionally, plant virus families detected in this study included *Solemoviridae*, *Alphaflexiviridae*, *Amalgaviridae*, *Betaflexiviridae*, *Bromoviridae*, *Closteroviridae*, *Endornaviridae*, *Luteoviridae*, *Tombusviridae*, and *Potyviridae*. As plant viruses are not known to infect humans, their detection in faecal samples most likely reflects the consumption of contaminated plant-derived foods or environmental exposure rather than active human infection (Zhang et al., 2006).

A diverse range of host-specific viruses not directly associated with gastroenteritis was also detected in the present study. These included fungi-infecting viruses belonging to the families *Botourmiaviridae*, *Botybirnaviridae*, *Chrysoviridae*, *Mitoviridae*, *Narnaviridae*, and *Totiviridae*. Avian-infecting viruses from the family *Circoviridae*, protist-infecting viruses from the family *Mimiviridae*, and insect-infecting viruses belonging to the family *Dicistroviridae* were also identified. The detection of these viruses highlights the remarkable diversity of the infant gut virome and suggests that they originate from multiple environmental, dietary, or microbial sources rather than representing viruses that actively infect humans.

Previous and recent studies have consistently shown that bacteriophages constitute most of the infant gut virome and are acquired from multiple sources, including the mother, breast milk, diet, and the environment (Fulci et al., 2021; Pannaraj et al., 2018; Breitbart et al., 2008). In contrast, bacteriophages accounted for only 6.1% of the total viral reads in the present study, with eukaryotic viruses predominating across all three cohorts. Among the bacteriophages, crAssphages were the most abundant, with the highest proportion observed in healthy controls, followed by malnourished children with gastroenteritis and non-malnourished children with gastroenteritis. Although crAssphages are regarded as the most abundant and ubiquitous bacteriophages in the human gut (Dutilh et al., 2014), they are typically acquired later in infancy, with their abundance increasing alongside that of their bacterial hosts (Liang et al., 2020; (Koenig et al., 2011). Our findings are consistent with those of Mogotsi et al., who also reported the presence of crAssphages in healthy South African children and similarly observed a predominance of eukaryotic viruses over bacteriophages (Mogotsi et al., 2024).

Members of the order *Caudovirales* were also detected across all study cohorts. Previous studies have suggested that the gut phageome may influence early-life growth and nutritional status, with specific bacteriophages being associated with growth impairment and stunting in children (Desai et al., 2020; Reyes et al., 2015). Although the source of bacteriophages could not be determined in the present study, they are known to be transmitted through several routes, including vertical transmission from mother to infant, faecal microbiota transplantation, and environmental exposure (Tamburini et al., 2018; Lim et al., 2015). Nevertheless, the role of bacteriophages in the pathogenesis of gastroenteritis and malnutrition remains poorly understood and warrants further investigation.

In the present study, mixed viral aetiology was observed among malnourished and non-malnourished children with gastroenteritis, as well as healthy controls, with frequent co-occurrence of multiple enteric viral genera, including *rotavirus* (RV), *norovirus* (NoV), *sapovirus* (SaV), *astrovirus* (AstV), *adenovirus* (AdV), *enterovirus* (EV), and *parechovirus* (PeV). This finding is consistent with previous reports showing that co-infections of multiple enteric pathogens is common among children with gastroenteritis (Nair et al., 2010). Similarly, a longitudinal study of healthy infants in the Free State Province, South Africa, reported frequent co-detection of gastroenteritis-associated viruses in the absence of clinical symptoms, with NoV–PeV, NoV–SaV, and SaV–PeV among the most common combinations (Milton T. Mogotsi et al., 2024). In the present study, RV was frequently involved in co-occurrence patterns and was detected in all children in the NDM control cohort. The detection of multiple enteric viruses in healthy children suggests that viral co-detection is not necessarily indicative of active disease and may reflect asymptomatic carriage, prolonged shedding, repeated exposure, or normal establishment of the infant gut virome (Carvalho et al., 2025; Mogotsi et al., 2024).

The relatively complex viral co-occurrence patterns observed among malnourished children with gastroenteritis may reflect alterations in the gut viral ecosystem associated with nutritional vulnerability. This is supported by Reyes et al. (2015), who reported disruption of normal gut virome development in Malawian children with severe acute malnutrition, including delayed maturation of bacteriophage and eukaryotic viral communities. Although, multiple concurrent enteric infections have been suggested to contribute to intestinal dysfunction, disease severity, malnutrition, and impaired growth in young children (Carvalho et al., 2025), mixed infections have also been associated with poor sanitation, inadequate hygiene, and limited access to safe water (Nimri et al., 2004). However, the cross-sectional design of the present study precludes establishing causal relationships between viral co-occurrence, malnutrition, and disease severity. Overall, the findings demonstrate that enteric viral co-occurrence is a common feature of the paediatric gut virome across nutritional and clinical groups, while the composition and complexity of viral combinations vary between cohorts.

This study has some limitations. The relatively small sample size and single-centre study design may limit the generalizability of the findings. In addition, the cross-sectional design and analysis of a single stool sample per participant precluded assessment of temporal changes in the gut virome and the duration of viral shedding. Although metagenomic next-generation sequencing enabled comprehensive detection of viral nucleic acids, it could not distinguish active infection from asymptomatic carriage, prolonged viral shedding, or transient viral passage. Furthermore, the limited availability of published data on the gut virome composition of malnourished children with gastroenteritis, particularly in South Africa, restricted direct comparison of the present findings with previous studies. This highlights the need for additional well-designed studies to strengthen the evidence base for understanding the role of the gut virome in childhood malnutrition and gastroenteritis.

## Conclusions

The application of mNGS enabled the first comprehensive characterisation of the faecal virome of malnourished South African children with gastroenteritis, providing valuable baseline data for an understudied paediatric population. The findings demonstrate that the infant gut virome is a complex and diverse ecosystem comprising human enteric and non-enteric eukaryotic viruses, as well as bacteriophages, across all study cohorts. Importantly, viral composition varied according to nutritional status. While *Rotavirus* predominated among healthy and non-malnourished children with gastroenteritis, Sapovirus was the dominant enteric virus among malnourished children with gastroenteritis. This finding suggests an association between nutritional status and virome composition that should be tested in larger, longitudinal studies. The frequent detection of enteric viruses among asymptomatic children further highlights the complexity of interpreting viral detection in early childhood, as viral carriage may reflect prolonged shedding, repeated exposure, or vaccine-derived viruses rather than active disease alone.

Overall, this study expands current knowledge of the paediatric gut virome in South Africa and demonstrates the complex viral landscape associated with childhood gastroenteritis irrespective of nutritional status. The detection of a broad range of established and potentially emerging enteric viruses underscores the importance of continued genomic surveillance alongside nutritional and preventive interventions to reduce the burden of gastroenteritis and malnutrition in young children. These findings also support continued surveillance and provide a rationale for longitudinal studies; they do not, by themselves, establish additional vaccine needs or identify vaccine targets. The baseline data generated provide a foundation for future longitudinal and comparative studies investigating the relationship between nutritional status, viral shedding, gut virome dynamics, and disease outcomes, with the goal of improving the prevention and management of childhood gastroenteritis and malnutrition in South Africa.

## Credit authorship contribution statement

Project Conceptualisation, Administration and Supervision:**, Kimesh L. Naidoo, Nathlee Shaik, Saheed Sabiu, and Martin M. Nyaga;** Methodology: **Marcelle M. Eyong, Milton T. Mogotsi, Ayodeji E. Ogunbayo, and Hlengiwe Sondlane**; Software: **Marcelle M. Eyong and Vivian C. Nwokorogu;** Sample collection and Data capture: **Nonkululeko G. Mabaso**; Formal analysis, Data curation, Validation: **Marcelle M. Eyong, Mathew D. Esona, and Vivian C. Nwokorogu;** Data visualisation: **Marcelle M. Eyong, and Vivian C. Nwokorogu**; Investigation: **Marcelle M. Eyong, Kimesh L. Naidoo, Nathlee Shaik, Saheed Sabiu, and Martin M. Nyaga;** Writing – original manuscript draft preparation: **Marcelle M. Eyong**; Writing-review and editing; **Marcelle M. Eyong, Milton T. Mogotsi, Ayodeji E. Ogunbayo, Hlengiwe Sondlane, Vivian C. Nwokorogu, Nonkululeko G. Mabaso, Mathew D. Esona, Kimesh L. Naidoo, Nathlee Shaik, Saheed Sabiu, and Martin M. Nyaga**; Funding acquisition: **Martin M. Nyaga**.

## Informed consent statement

Written informed consent was obtained from the parents and/or legal guardians of all participating children prior to sample collection.

## Ethical statement

Ethical approval for this study was obtained from the Biomedical Research Ethics Committee (BREC) of the University of KwaZulu-Natal under ethics number BREC/000040001/2022, which was renewed on 20/03/2025 by the KwaZulu-Natal Department of Health (KZ_202112_002). Approval was also obtained from and the management of King Edward VIII Hospital (KE 2/7/16 (11/2022). Additional approval was granted by the Health Sciences Research Ethics Committee (HSREC) under ethics number UFS-HSD2024/2670 and by the Environment and Biosafety Research Ethics Committee (EBREC), ethics number UFS-ESD2024/0440 of the University of the Free State in Bloemfontein, South Africa.

## Acknowledgments

The authors gratefully acknowledge the partial tuition bursary awarded to Ms. Marcelle Eyong Manyi by the Centre for Graduate Support through the University of the Free State 2026 Postgraduate Bursary Programme. We sincerely thank all the infants and their parents or legal guardians for participating in this study. We also acknowledge the management of Victoria Mxenge Hospital, Durban, and the staff of the Paediatric Department for granting permission to conduct this study and for their invaluable support during participant recruitment, data collection, and faecal sample collection. We are especially grateful to Mrs. Leora Sewnarain for her assistance in obtaining the administrative approvals required for this study. We further thank Nkosazana D. Shange, Surprise Baloyi, Angela Mbele, Thabisa Mpaxa, and Mbali Faith Ncube for their assistance with the laboratory experiment.

## Supplementary Information

Supplementary material will be made available online.

## Data availability

The data supporting the findings of this study are available within the article and its Supplementary Information. The raw sequencing data have been deposited in the National Centre for Biotechnology Information (NCBI) Sequence Read Archive (SRA) under BioProject accession number **PRJNA1475839**.

